# Longitudinal accumulation of *in vivo* and *in vitro-*grown *Treponema pallidum* subsp. *pallidum* TprK variants in the presence and absence of immune pressure

**DOI:** 10.1101/2021.06.26.450029

**Authors:** Michelle J. Lin, Austin M. Haynes, Amin Addetia, Nicole A.P. Lieberman, Quynh Phung, Hong Xie, Tien V. Nguyen, Barbara J. Molini, Sheila A. Lukehart, Lorenzo Giacani, Alexander L. Greninger

## Abstract

Immune evasion by *Treponema pallidum* subspecies *pallidum* (*T. pallidum*) has been attributed to antigenic variation of its putative outer-membrane protein TprK. In TprK, amino acid diversity is confined to seven variable (V) regions, and generation of sequence diversity within the V regions occurs via a non-reciprocal segmental gene conversion mechanism where donor cassettes recombine into the *tprK* expression site. Although previous studies have shown the significant role of immune selection in driving accumulation of TprK variants, the contribution of baseline gene conversion activity to variant diversity is less clear. Here, combining longitudinal *tprK* deep sequencing of near clonal Chicago C from immunocompetent and immunosuppressed rabbits along with the newly developed *in vitro* cultivation system for *T. pallidum*, we directly characterized TprK alleles in the presence and absence of immune selection. Our data confirm significantly greater sequence diversity over time within the V6 region during syphilis infection in immunocompetent rabbits compared to immunosuppressed rabbits, consistent with previous studies on the role of TprK in evasion of the host immune response. Compared to strains grown in immunocompetent rabbits, strains passaged *in vitro* displayed low level changes in allele frequencies of TprK variable region sequences similar to that of strains passaged in immunosuppressed rabbits. Notably, we found significantly increased rates of V6 allele generation relative to other variable regions in *in vitro* cultivated *T, pallidum* strains, illustrating that the diversity within these hypervariable regions occurs in the complete absence of immune selection. Together, our results demonstrate antigenic variation in *T. pallidum* can be studied *in vitro* and occurs even in the complete absence of immune pressure, allowing the *T. pallidum* population to continuously evade the immune system of the infected host.

**Author Summary:** Syphilis continues to be a disease of global and public health concern, even though the infection can be easily diagnosed and effectively treated with penicillin. Although infected individuals often develop a strong immunity to the pathogen, repeated infection with the syphilis agent, *Treponema pallidum* subspecies *pallidum* (*T. pallidum*), is possible. Several studies point at antigenic variation of the *T. pallidum* TprK protein as the mechanism responsible for evasion of the immunity that develops during infection, pathogen persistence, and re-infection. Past studies have highlighted the importance of immune clearance of dominant variants that, in turn, allows less represented variants to emerge. The contribution of an immunity-independent baseline generation of variability in the *tprK* gene is less clear. Here, we used deep sequencing to profile *tprK* variants using a laboratory-isolated *T. pallidum* strain nearly isogenic for *tprK* that was propagated over time *in vitro*, where no immune pressure is exerted on the pathogen, as well as in samples obtained from immunosuppressed and immunocompetent rabbits infected with the same strain. We confirmed that *tprK* accumulates significantly more diversity under immune pressure, and demonstrated a low but discernible basal rate of gene conversion in complete absence of immune pressure.

## Introduction

Syphilis, caused by the spirochete *Treponema pallidum* subspecies *pallidum* (*T. pallidum*), remains a public health priority with its estimated global burden of 36 million infections and steadily increasing incidence in high-income countries, particularly in men who have sex with men (MSM) and persons living with HIV [1–6]. Evidence that syphilis rates continue to rise despite the availability of inexpensive and effective treatment highlights the need for greater understanding of syphilis pathogenesis to improve disease control measures and foster development of an effective syphilis vaccine. The feasibility of this work, however, has been hampered by many factors, such as our historical inability to cultivate *T. pallidum in vitro* and the inability to genetically engineer this pathogen, which have only recently been overcome [7, 8].

Early syphilis infection manifests in distinct clinical stages, followed by a period of latency that, in most cases, will last for the remainder of the patient’s lifetime, even in absence of treatment [9]. Data from the pre-antibiotic era, however, showed that in about 30% of the patients, the disease reactivates after years to decades from the initial exposure to *T. pallidum*, and progresses to its tertiary stage, with serious clinical manifestations. The astounding, decade-long persistence of *T. pallidum* in the host is likely the result of a multiplicity of strategies evolved by this pathogen to become an effective obligated human pathogen. Among these strategies, pivotal is *T. pallidum* ability to actively evade the host immune response during infection, currently attributed to antigenic variation of the putative outer membrane protein TprK [10–12]. TprK sequence diversity is generated by non-reciprocal segmented gene conversion of 53 donor sites within seven discrete variable regions (named V1 – V7) dispersed throughout the gene length [13–15]. A previous study demonstrated that *tprK* sequence variability accumulates more rapidly in presence of immune pressure, particularly in V6 [16]. This study, along with others [17, 18], clearly showed the role of the host immune selection in the process of accumulation of *tprK* variants in an immunocompetent experimental subject.

The above results, however, like most of those that improved our molecular understanding of *T. pallidum* pathogenesis, are based on samples obtained from experimentally infected rabbits [19–22], due to the inability of *T. pallidum* to survive outside the mammalian host [23]. This limitation hindered our ability, for instance, to understand the dynamics of *tprK* variant generation in an environment completely devoid of immune pressure on the pathogen, and where the pathogen cannot freely disseminate from the site of infection. Recently however Edmondson *et al.* [7] developed a tissue culture system able to stably propagate *T. pallidum in vitro* by perfecting a cell culture-based system initially pioneered by Fieldsteel *et al.* [24]. Here, to expand on these previous studies of *tprK*, we paired *in vitro* cultivation and deep sequencing to profile the evolution of *tprK* diversity in cultured laboratory-isolated Chicago C *T. pallidum* cells nearly isogenic for *tprK.* We compared diversity accumulation in the *in vitro*-passaged Chicago C to experimental infection by Chicago C in immunocompetent and immunosuppressed animals. Our results confirm significantly greater antigenic variation, particularly in V6, during syphilis infection in both immunocompetent vs. immunosuppressed rabbits, and in immunocompetent rabbits vs. culture system. We also demonstrate higher rates of V6 diversity generation relative to other variable regions in both immunosuppressed rabbits and in *in vitro*-propagated treponemes, suggesting that a low level basal rate of gene conversion occurs in absence of immune pressure.

## Methods

### Ethics statement

For the immunosuppressed vs. immunocompetent rabbit studies, *T. pallidum* Chicago strain was propagated in New Zealand white rabbits as previously described [16]. For the *T. pallidum in vitro* culture vs. control rabbit studies, male New Zealand white rabbits were housed at the University of Washington (UW) Animal Research and Care Facility (ARCF). Care was provided according to the procedures outlined in the Guide for the Care and Use of Laboratory Animals under the protocols approved by the UW Institutional Animal Care and Use Committee (IACUC; Protocol # 4243-01, PI: Lorenzo Giacani). Specific pathogen free (SPF; *Pasteurella multocida* and *Treponema paraluiscuniculi*) animals, weighing approximately 3.5-4.5 kilograms were purchased from the Western Oregon Rabbit Company (Philomath, OR). Upon arrival at the ARCF, rabbits were allowed to acclimate prior to initial serological testing to confirm lack of *T. paraluiscuniculi* infection, a precaution taken because only a random subset of animals are tested by the vendor. All animals were tested using the Venereal Disease Research Laboratory (VDRL, Becton Dickenson, NJ) and Fluorescent treponemal antibody-absorption (FTA-ABS, Trinity Biotech, Ireland) tests. Both tests were performed according to manufacturer’s instructions aside from the substitution of a secondary FITC-labelled goat anti-rabbit IgG rather than the anti-human secondary antibody included in the FTA-ABS kit. Upon confirmation of negative serological reactivity, rabbits were cleared for use in these studies.

### Experimental infection of immunocompetent vs. immunosuppressed rabbits

In a previously published study [16], five pharmacologically immunosuppressed rabbits and five immunocompetent control rabbits were infected intradermally with the Chicago C *T. pallidum* strain that had been previously derived through *in vivo* cloning and was known to be nearly isogenic at the *tprK* locus. Briefly, rabbits were injected at 10 sites each on their clipped backs and 10^6^ treponemes were injected per site. Pharmacological immunosuppression was achieved by weekly intramuscular (IM) injections of 20 mg methylprednisolone acetate (Sicor, Irvine, CA). Treatment was started 3 days before experimental infection, and doses were administrated weekly for a total of six injections. A lesion biopsy (4-mm punch biopsies taken under local lidocaine anesthesia) was harvested from each rabbit weekly for a period of 5 weeks; these samples were used for DNA extraction and *tprK* profiling in the present study as shown in Fig 1A schematic.

**Fig 1.**
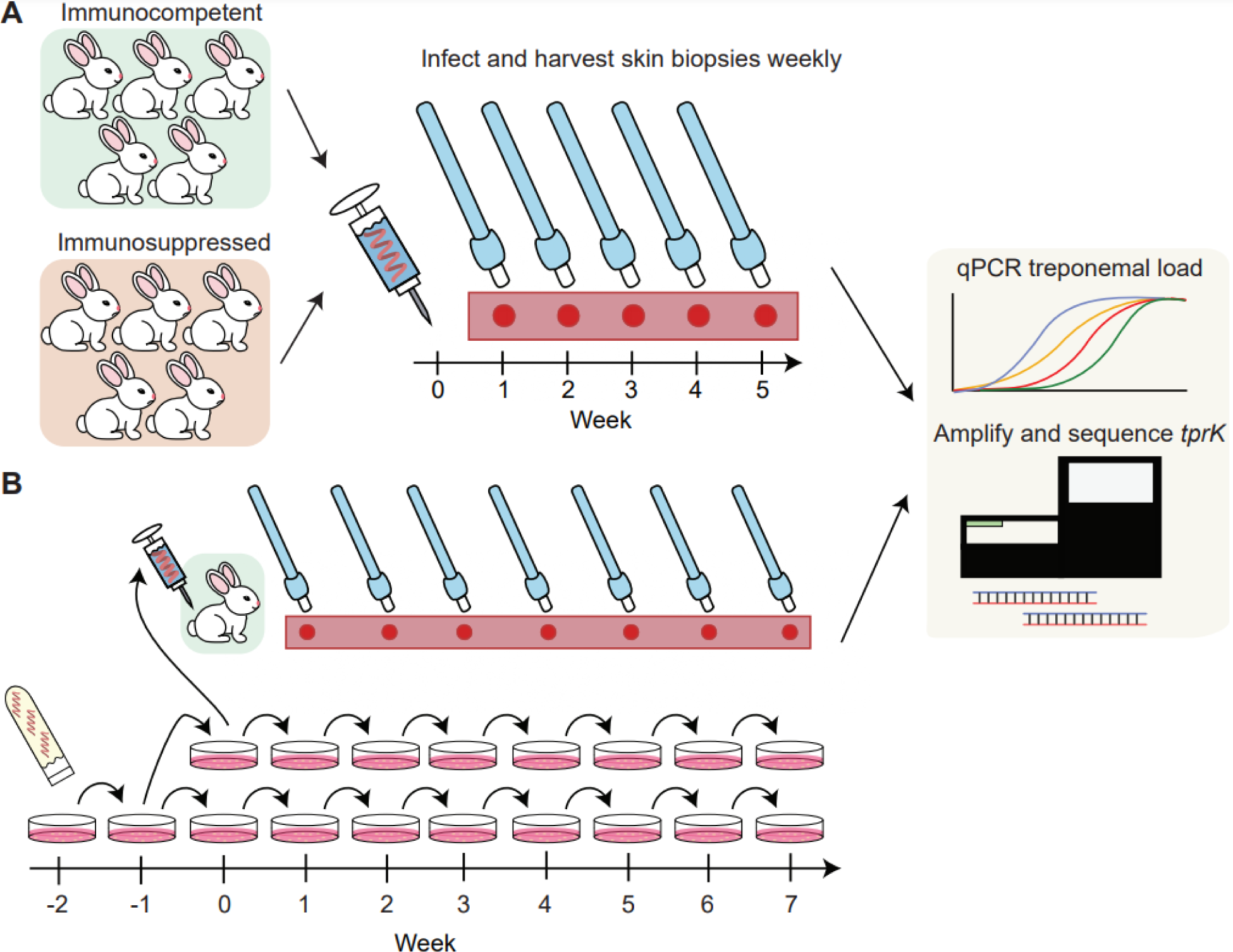
Study design for sample acquisition from (A) immunosuppressed vs. immunocompetent rabbits and (B) *in vitro* propagated treponemes vs. control. (A) Five immunocompetent rabbits and five immunosuppressed rabbits were infected with inoculum at Week 0. Harvesting of punch biopsies from each rabbit was done weekly for 5 weeks. Samples were analyzed for treponemal load by qPCR and *tprK* sequence by next- generation sequencing. (B) Glycerol stock of *T. pallidum* Chicago C was thawed at Week -2, passaged *in vitro* at Week -1 and again at Week 0, at which point treponemes were 1) inoculated intradermally into a control rabbit, and 2) seeded into two sets of weekly-passaged cultured samples for 7 weeks. Samples were collected weekly from punch biopsies of the rabbit and from cell culture, then analyzed for treponemal load and *tprK* sequence.

### Parallel in vivo and in vitro cultivation of T. pallidum

One day prior to *in vitro* inoculation of the Chicago C strain of *T. pallidum*, 10^5^ rabbit Sf1 epithelial cells were plated into each well of a 6-well plate and incubated at 37°C overnight in a 5% CO2 atmosphere within a HeraCell 150 incubator (Thermo Fisher Scientific, Waltham, MA). At the same time, treponemal culture medium (TpCM2) was prepared according to the Edmondson *et al.* protocol [25], and incubated overnight at 34°C in a microaerophilic atmosphere (1.5% O_2_, 5% CO_2_ and 93.5% N_2_) to deplete the medium of oxygen.

The subsequent day, Sf1 epithelial cells were rinsed with equilibrated TpCM2 medium and then equilibrated in 5 mL of media in the microaerophilic incubator for three hours. A frozen glycerol stock of *T. pallidum* strain Chicago C was thawed at room temperature and bacteria were enumerated on a Leica DM2500 darkfield microscope (Leica, Wetzlar, Germany) to inoculate 10^6^ bacteria into each well of the 6-well plate. The plate was then returned to the microaerophilic incubator for seven days. Residual inoculum bacteria were collected via μL of buffer ATL (Qiagen, Hilden, Germany) for DNA extraction. After seven days, the culture was retrieved, and supernatants were discarded. Treponemes were then released from the epithelial cell layer via trypsin digestion, enumerated as described above and passaged (Figure 1B). Starting with the week 0 culture, inocula were increased to 10^7^ cells per well and the bacteria were seeded into two 6-well culture dishes. At every passage starting from Week 0, cultured treponemes were harvested from matching wells from both plates and pelleted via centrifugation. The remaining wells were used to seed two additional plates. In parallel, the same treponemes in Week 0 were used to infect a New Zealand white rabbit. Prior to rabbit infection, treponemes were enumerated once more on a Nikon OptiPhot2 darkfield microscope (Nikon, Tokyo, Japan) to ensure viability and correct density. After verifying viability, 10^6^ bacteria per site were administered intradermally at eight sites on the shaved back of the rabbit. The rabbit was monitored briefly and then returned to its cage. After this point, infected rabbits were shaved daily to allow monitoring of lesion development and facilitate weekly biopsy collection. Starting a week after infection (Week 1), a single lesion developing on the back of the rabbit was harvested using a 4-mm dermal biopsy punch. Tissue biopsies harvested from week 1 to week 7 were minced using sterile scalpels μL of lysis buffer for DNA extraction.

### DNA extraction and real-time quantification of treponemal load

Genomic DNA was extracted using the QiaAmp DNA Mini Kit (Qiagen, Valencia, CA) μL of sample suspension per manufacturer’s protocol. DNA was then eluted into a 100 μL were determined by qPCR using forward primer 5’- CAA GTA CGA GGG GAA CAT CGA T-3’, reverse primer 5’-TGA TCG CTG ACA AGC TTA GG-3’, and AmpliTaq Gold 360 spiked with 1x SYBR Green. Amplification conditions for qPCR were as follows: 95°C for 10 minutes, followed by 45 cycles of 95°C for 30 seconds, 56°C for 30 seconds, 72°C for 30 seconds.

### Quantification of rabbit CFTR DNA

Levels of CFTR DNA were determined using a droplet digital PCR (ddPCR) assay on the QX100 ddPCR system (Bio-Rad Laboratories, Hercules, CA), using CFTR sense (5’-GCG ATC TGT GAG TCG AGT CTT-3’) and CFTR anti-sense (5’-GGC CAG ACG TCA TCT TTC TT-3’) primers and probe (6FAM-CCAAATCCATCAAACCATCC-NFQMGB).

The ddPCR reaction mixture consisted of 12.5 μL of primers and probe mix to a final concentration of 900nM primers and 250nM probe, and 1 μ L. Reaction mixture was then loaded onto Automated Droplet Generator (Bio-Rad Laboratories, Hercules, CA). The generated droplets were subsequently transferred to a 96-well PCR plate and PCR amplification was performed on a C1000 Touch Thermal Cycler (Bio-Rad) with the following conditions: 95°C for 10 minutes, 40 cycles of 94°C for 30 seconds and 60°C for 1 minute, followed by 98°C for 10 minutes and ending at 4°C. After amplification, the plate was loaded onto the droplet reader (Bio-Rad). Data was analyzed with QuantaSoft analysis software and quantitation of target molecules presented μL of PCR reaction.

### TprK library preparation and next-generation sequencing

TprK libraries were prepared similarly as previously described [15]. Thirty-five cycles of L reaction with 0.4 mM barcoded TprK forward and reverse primers (S1 Table) using the 2x CloneAmp MasterMix (Takara) with a 62°C annealing temperature. For the immunocompetent versus immunosuppressed rabbit experiment, the μL) of genomic DNA was used for PCR, including a median of 73681 (range: 140 to 6269883) genome copies. For the culture versus immunocompetent rabbit experiment, a range between 3379 - 211944 (median: 143036) genome copies were used, with the number of input genomes for each timepoint normalized to the rabbit sample collected that week. PCR products were run on a 1% agarose gel to confirm the presence of a band at approximately 1.6 kb and cleaned with 0.6x Ampure beads (Beckman Coulter). Two nanograms of PCR product was used in a two-fifths Nextera (Illumina) library prep with 15 cycles of amplification. Low molecular weight products were removed with 0.6x Ampure beads, yielding final libraries with a minimum size of approximately 300 bp. Libraries were quantified, pooled, and run on a 1x192 single end run on an Illumina Miseq. A second replicate of each sample was prepared as described above to control for potential polymerase error. Samples yielding a minimum of 1000 mean coverage mapping to *tprK* were kept for downstream analysis, with raw reads ranging from 15461 – 368148 reads.

### Sequencing analysis

*tprK* reads were analyzed with custom python and R scripts. Briefly, raw reads were adapter-trimmed and filtered for quality scores above Q20 using trimmomatic v0.39 [26].

Variable regions were obtained from all samples by fuzzy matching of the flanking conserved sequences, allowing for a maximum of 3 mismatches. To reduce low-level polymerase errors, only high confidence sequences that were above a relative frequency of 0.1% in both Illumina library preparations were included in our analyses. We used the R package VEGAN [27] to calculate Pielou’s evenness and Shannon diversity measures for each sample.

### Donor site recombination

We constructed recombination possibilities of our *tprK* alleles according to the previously characterized *tprK* anatomy of donor site segments separated by occasional overlapping 4-bp internal repeats [13]. We used blastn to extract exact matches greater than or equal to 5 nucleotides in length within the 12.5-kb locus containing the *tprD* gene, using only matches with at least 10 reads of support in any given sample. We then used these segments to recursively build on each other to recreate each unique *tprK* allele, allowing exact or overlapping 1-2 internal repeats between segments. We allowed for up to 5% of no coverage in each variant in order to account for the possibility of segments that are <4 bp long — below the minimum length requirement for blastn — and for differences in our variable region boundary definitions compared to what was seen when *tprK* anatomy was previously characterized [13,15,28].

### Data Availability

Reads from *tprK* sequencing of the samples used in this study are available under the NCBI BioProject number PRJNA734645. All code used for analysis is publicly available on GitHub (https://github.com/greninger-lab/longitudinal_tprk).

## Results

### Longitudinal analysis of tprK in multiple model systems

We profiled longitudinal evolution of *tprK* in treponemes obtained from multiple model systems over two experiments. In the first experiment, longitudinal samples were obtained from skin lesions of five pharmacologically immunosuppressed rabbits and five immunocompetent rabbits infected with the *tprK*-isogenic Chicago C strain (Fig 1A). In the second experiment, we propagated the Chicago C strain in culture and also infected one additional control animal, harvesting samples from each source every week for seven weeks after initial inoculation (Fig 1B).

The quantity of *T. pallidum* in each sample at the time of harvest was determined using quantitative PCR targeting the *T. pallidum tp0574* gene. A median *tp0574*/CFTR copy number ratio of 0.64 was measured in the immunosuppressed vs. immunocompetent rabbit experiment, while the median copy number ratio was 6.75 the *in vivo* vs. *in vitro* experiment, mostly due to treponemal enrichment in the *in vitro* culture. As expected, there was a noticeable but non-significant decrease in copy number over time in immunocompetent rabbits compared to immunosuppressed rabbits (Fig 2A, S1 Fig) due to immune clearance of *T. pallidum* occurring during early infection. This is consistent with previous results and likely contributed to the decrease in diversity observed in later weeks in certain variable regions. By week five, all samples collected from immunocompetent rabbits in the first experiment had less than 5.4 x 10^-3^ *tp0574*/CFTR copy number ratio. In contrast, at week 7 of the second experiment, all samples harvested from the immunocompetent rabbit had greater than 3.0 *tp47*/CFTR copy number ratio (Fig 2B).

**Fig 2.**
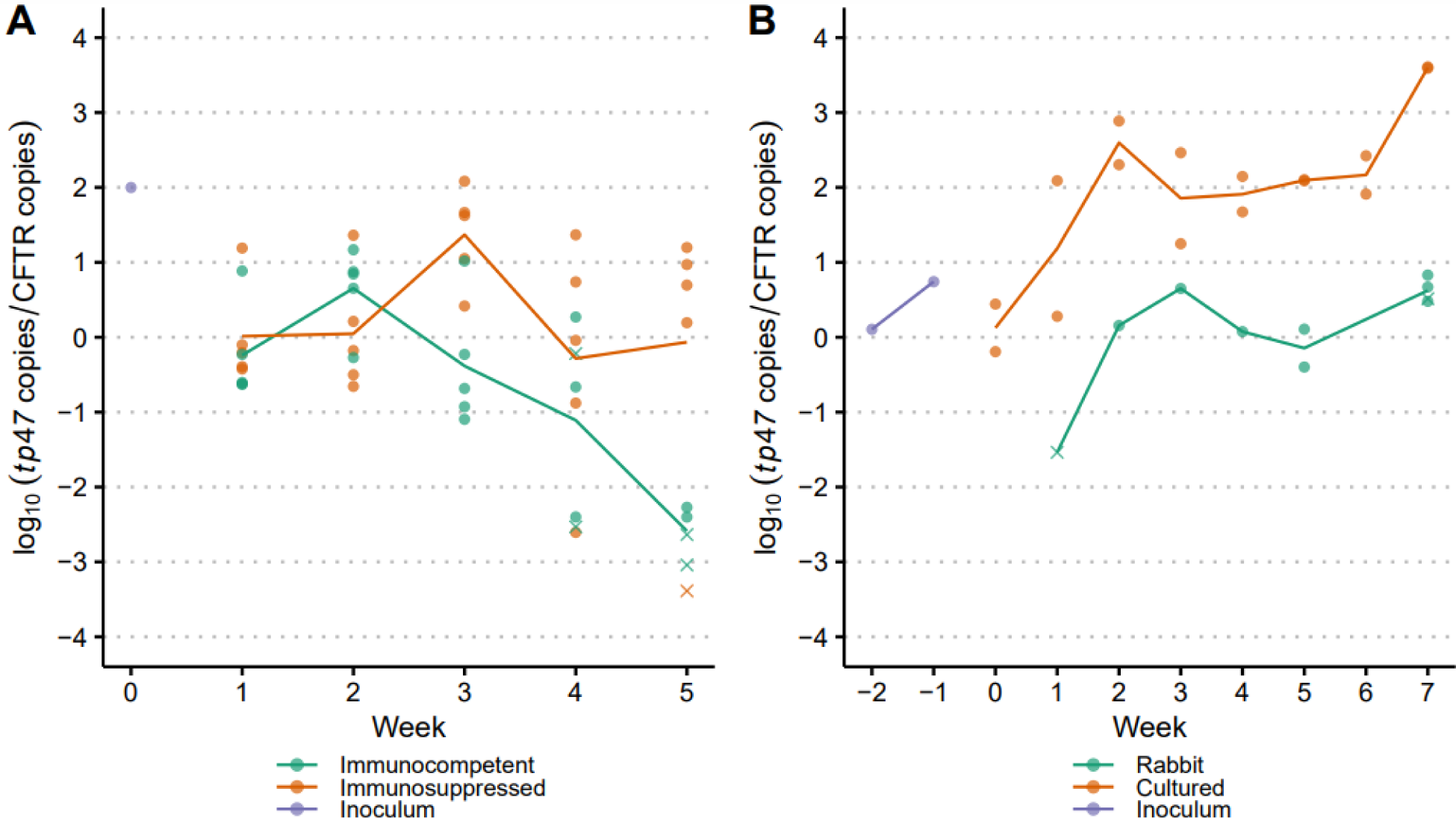
Log copy numbers for samples passaged in (A) immunosuppressed vs. immunocompetent rabbits and (B) immunocompetent rabbit vs. *in vitro* culture system. Each unique sample is represented by either a dot or a cross, where a cross indicates that we were unable to successfully recover sequence from the sample. Lines show the mean normalized treponemal load, as quantified by the log ratio of *tp0574* copies to CFTR copies and are grouped and color-coded by passage type.

We then amplified the *tprK* gene and performed next-generation sequencing on samples with recoverable *tprK* PCR product. Sequencing data for all seven variable regions at varying timepoints were successfully obtained for 23 immunosuppressed rabbit samples and 19 immunocompetent rabbit samples (Fig 2A), as well as 8 samples from the control rabbit, 8 samples each from the two replicate culture sets, and 2 early passage culture specimens (labeled “Inoculum”, Fig 2B).

To examine *tprK* heterogeneity, we extracted high-confidence *tprK* alleles for each variable region above a relative frequency of 0.1% in two Illumina library preparations for further analysis [15]. Sequences were reproducible (r^2^ = 0.997) across the two technical replicates for all samples (S2 Fig). Individual longitudinal plots for each sample with their unique *tprK* sequences are available for both experiments in S3 Fig and S4 Fig respectively. In all model systems across all timepoints, we identified the least number of unique sequences in V4 (median: 5, range: 1 – 9) and the most in V6 (median: 44, range 5 – 89). Longitudinally across the 5-week period of the immunocompetent vs. immunosuppressed rabbit experiment, the number of unique V region alleles inclusive of all variable regions increases from a mean of 3.0 sequences in the original inoculum (range: 2 – 5) to a mean of 15.0 sequences in the last sampled week (range: 1 – 58) (S2 Table). Similarly, across the 10-week period of the *in vivo* vs. *in vitro* experiment, the number of unique V region alleles increases from the original inoculum (mean: 9.28, range: 3 – 18) to last passaged week (mean: 23.3, range: 6 – 89). In addition, we observed decreased clonality of the highest frequency sequence over time as lower frequency variants become more prevalent in immunocompetent rabbits, compared to samples passaged in immunosuppressed rabbits or in culture.

### Longitudinal TprK evolution is significantly greater in immunocompetent rabbits versus both immunosuppressed rabbits and in vitro culture

Next, we analyzed the accumulation of diversity in each variable region for all samples in both experiments. A prior investigation examined differences in accumulation of diversity between these same immunocompetent versus immunosuppressed rabbits through fragment length analysis, with values of the reciprocal of Simpson’s diversity index (RSDI) differing significantly (p<0.05) at 3 weeks after infection for V6 [16]. Here, we used two library preparations to call high-confidence sequences at depths that allowed better capture of the true diversity of *tprK* variants. An additional metric, Pielou’s evenness, was used to quantify diversity as the extent of equitability in the distribution of observed variants in each variable region. We confirmed a significant difference in Pielou’s evenness score in V6 of immunocompetent rabbits versus immunosuppressed rabbits beginning at Week 3 (p<0.0001) (Fig 3A, S2 Table), but we did not observe any significant difference in mean diversity between these two groups of rabbits in any other variable region.

**Fig 3.**
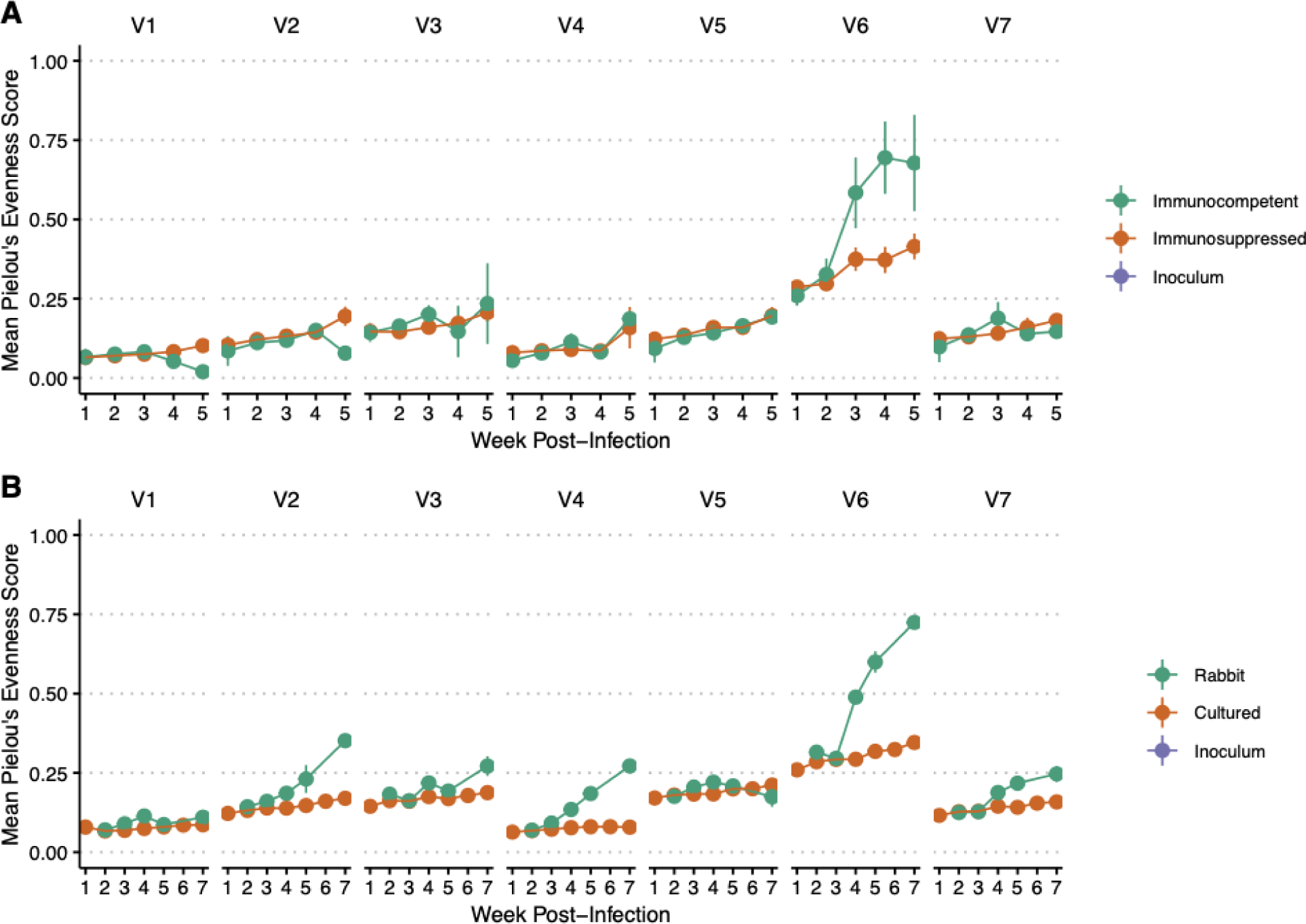
Longitudinal allelic diversity of samples passaged in (A) immunosuppressed vs. immunocompetent rabbits and (B) immunocompetent rabbit vs. *in vitro* culture system. Data are separated by variable region. Dots represent the mean Pielou’s evenness score across different color-coded groups, with lines marking the standard deviation.

We then assessed the longitudinal TprK evolution in the novel culture system by comparing *tprK* sequences from samples passaged in a single immunocompetent rabbit and samples passaged *in vitro*. Similar to the results from immunocompetent vs. immunosuppressed rabbits, we observed a 1.7-fold increase in Pielou’s evenness score of V6 in samples passaged *in vivo* compared to *in vitro* starting at week four post-infection (Fig 3B). Using an unpaired two- sample t-test, we also found a significantly greater number of unique *tprK* V6 alleles in the control rabbit than cultured samples (p=0.015). In contrast to the previous experiment, using deep sequencing, we additionally observed a significant accumulation of variable region diversity over time in V2, V3, V4, and V7 in the control rabbit compared to *in vitro* samples. Overall, these data demonstrate a significant increase of TprK variants, especially in V6, in immunocompetent rabbits compared to both immunosuppressed rabbits and *in vitro* culture, confirming that immune selection plays a significant role in the appearance of TprK variants.

### Accumulation of unique V6 alleles occurs even without immune selection and is reproducible across culture replicates in all variable regions

We further investigated accumulation of V region alleles in immunosuppressed rabbits and in culture, where variants were generated at similar rates (Fig 4, Table 1). Interestingly, in both types of passaging, V6 generated significantly more unique variants over time than in the other variable regions (p<0.0001), suggesting that greater V6 accumulation of new variants is not solely a function of immune selection (Table 1). To estimate the baseline rate of sequence change, we examined the weekly decrease of the most common sequence for each variable region because of the extremely low frequency levels of the many newly generated sequences.

**Fig 4.**
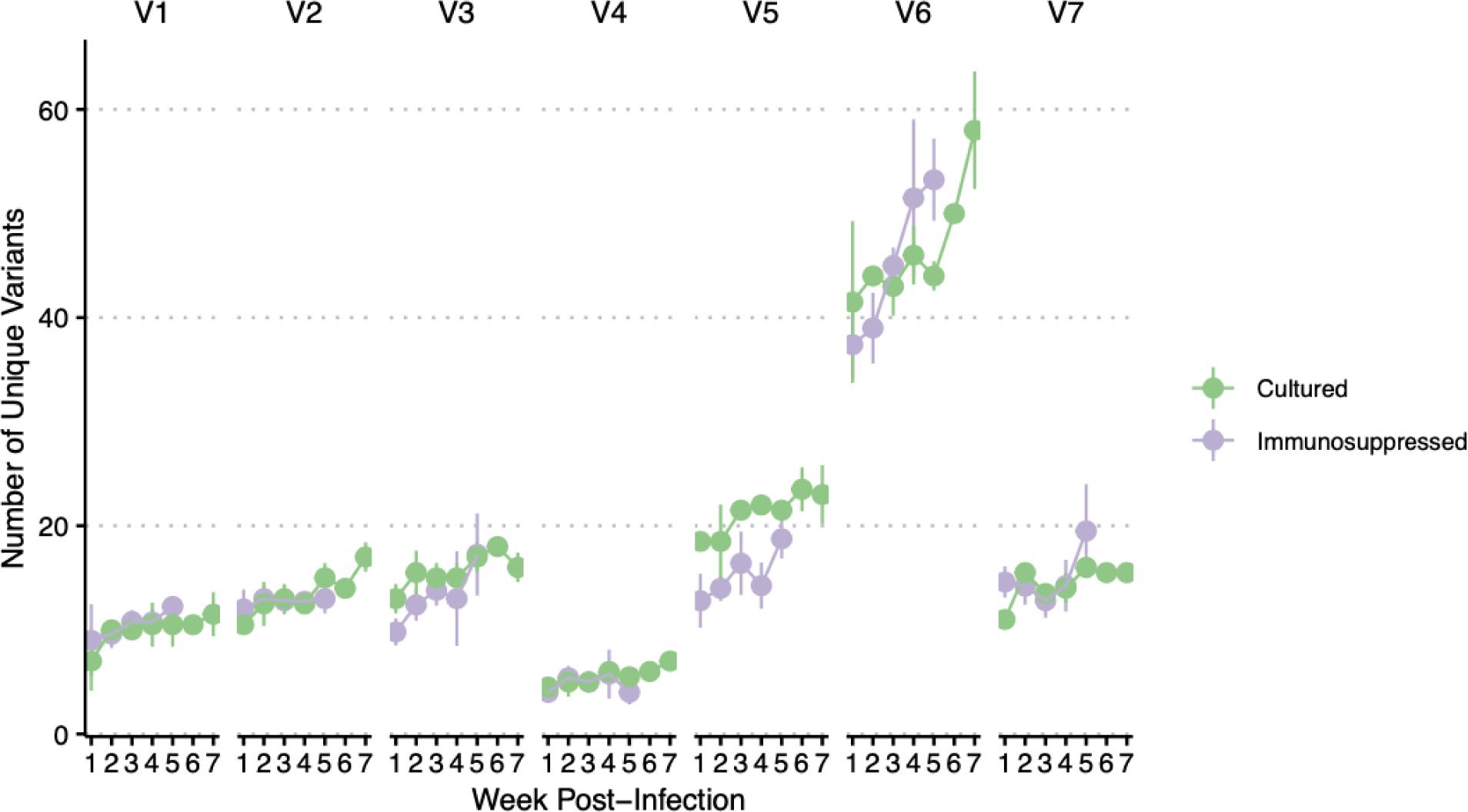
Comparison of number of unique V region alleles detected in samples passaged in immunosuppressed rabbits and in culture. X-axis depicts weeks post-infection. Dots represent the mean number of unique variants at that time point, with lines demonstrating standard deviation.

**Table 1.**
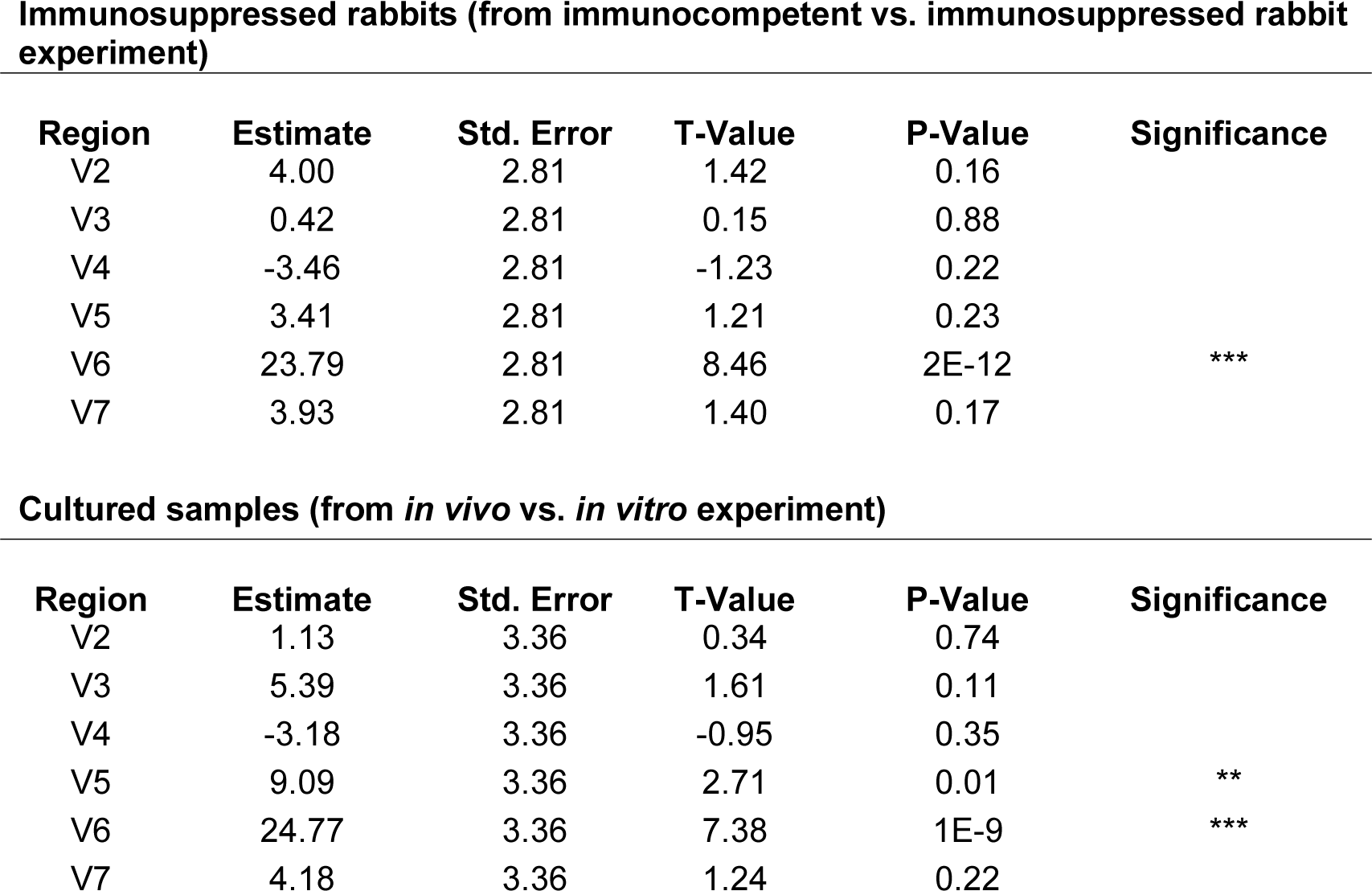
Linear mixed model shows significant increase in unique variants in V6 in both immunosuppressed rabbits and cultured samples. Change in unique variants in each variable region is compared to V1 for both experiments using T-Tests following Kenward- Roger’s method [29]. Significance is denoted with asterisks (0***, 0.001**, 0.01*).

**Immunosuppressed rabbits (from immunocompetent vs. immunosuppressed rabbit experiment)**
| Region | Estimate | Std. Error | T-Value | P-Value | Significance |
| --- | --- | --- | --- | --- | --- |
| V2 | 4.00 | 2.81 | 1.42 | 0.16 |  |
| V3 | 0.42 | 2.81 | 0.15 | 0.88 |  |
| V4 | -3.46 | 2.81 | -1.23 | 0.22 |  |
| V5 | 3.41 | 2.81 | 1.21 | 0.23 |  |
| V6 | 23.79 | 2.81 | 8.46 | 2E-12 | *** |
| V7 | 3.93 | 2.81 | 1.40 | 0.17 |  |

**Table 1. Linear mixed model shows significant increase in unique variants in V6 in both immunosuppressed rabbits and cultured samples.**
| Region | Estimate | Std. Error | T-Value | P-Value | Significance |
| --- | --- | --- | --- | --- | --- |
| V2 | 1.13 | 3.36 | 0.34 | 0.74 |  |
| V3 | 5.39 | 3.36 | 1.61 | 0.11 |  |
| V4 | -3.18 | 3.36 | -0.95 | 0.35 |  |
| V5 | 9.09 | 3.36 | 2.71 | 0.01 | ** |
| V6 | 24.77 | 3.36 | 7.38 | 1E-9 | *** |
| V7 | 4.18 | 3.36 | 1.24 | 0.22 |  |

We observed a mean weekly decrease of 2.40% relative frequency in the most highly abundant sequence in V6 in the immunosuppressed rabbits compared to a mean weekly decrease ranging between 0.37% and 1.04% for the other variable regions (S3 Table). In the cultured treponemes, we similarly observed a mean weekly decrease of 2.25% in the most highly abundant sequence in V6 compared to a mean weekly decrease ranging between 0.16% and 0.71%.

To better understand how these variants are being generated over time, we compared the generation of V region sequences between our two sets of samples propagated in culture. In a purely stochastic model, we would expect to see emergence of several sequences present in one set that are absent in the other. However, we observed that the relative frequencies of *tprK* alleles were remarkably reproducible across our two culture replicates for all variable regions (r^2^ = 0.9998) (Fig 5A). We did not observe any variable region sequences above 0.62% relative frequency present in one culture replicate that was not present in the other (Fig 5B). In addition, there were a total of 128 unique alleles over all variable regions generated in culture that were not present in the inoculum above a relative frequency of 0.1% in both library preparations.

**Fig 5.**
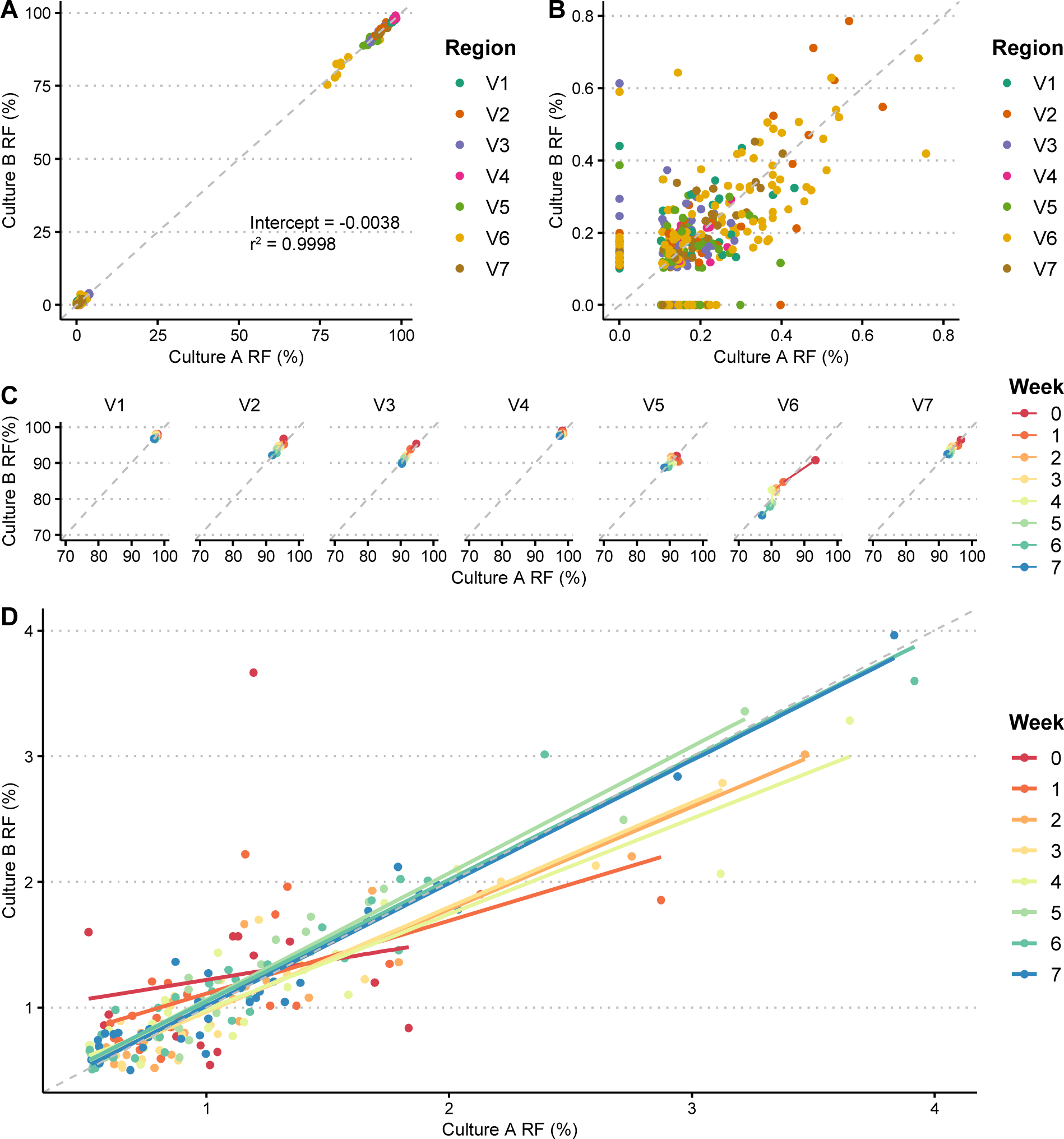
Similar relative frequencies (RFs) of *tprK* variable region alleles are observed across culture replicates. (A) Comparison of relative frequencies of alleles between two sets of culture passages. Dots represent unique variable region sequences of varying timepoints, color-coded by region. Dotted gray line plots Y=X. (B) Comparison of relative frequencies of all novel alleles not in inoculum between two culture replicates, color-coded by variable region. (C) Relative frequencies of alleles between both culture replicates, at over 70%. Data are grouped by variable region and color coded by week. (D) Correlation between relative frequencies of alleles in culture replicates at low frequencies over time. Dots represent unique variable region sequences between timepoints, with corresponding regression lines plotted. Both are color- coded by week.

These novel alleles stayed at low frequencies, reaching a maximum of 0.79% relative frequency (Fig 5B). Predictably, at high relative frequencies (70%), the clonal sequence decreased in frequency from a mean of 95.46% to 90.67% across all variable regions over time (Fig 5C) as lower-level V region alleles increased in number. At lower relative frequencies (0.5 – 4%), the pattern became less clear, though over time relative frequencies of alleles in both culture replicates trended towards closer shared identity (Fig 5D and Fig 6), and often increased in frequency together.

**Fig 6.**
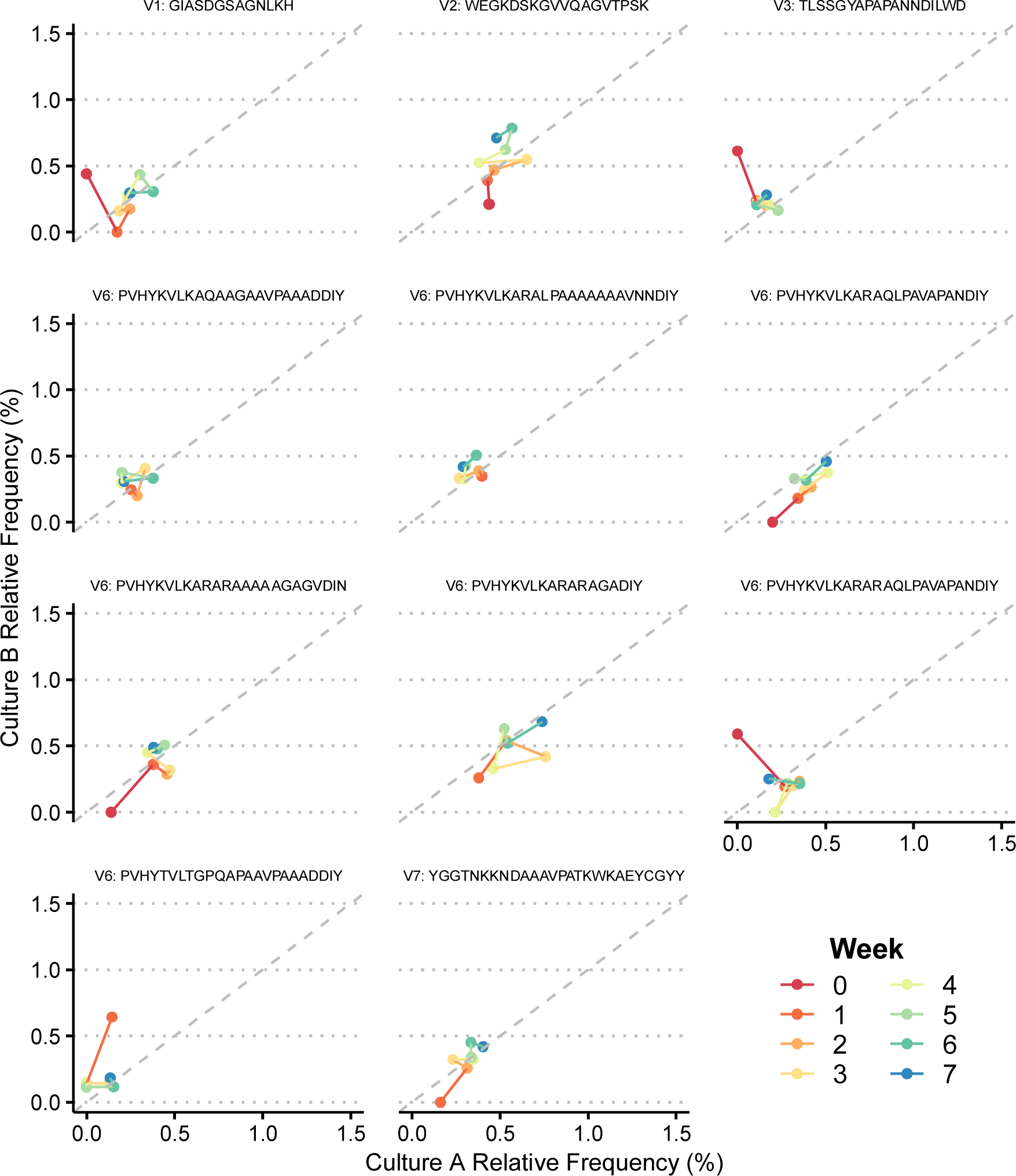
Comparison between culture replicates of individual novel V region alleles absent in inoculum that reached ≥0.4% relative frequency at any timepoint. Unique alleles are labeled above each plot with their variable region and amino acid sequence. Lines sequentially connect each timepoint and are color-coded by week.

### T. pallidum employs multiple strategies to generate antigenic variation via donor site recombination

We further examined how *tprK* sequences were being constructed *in vitro* through recombination of donor sites according to the TprK sequence anatomy previously characterized by Centurion-Lara *et al*. [11]. In this model, donor site segments surrounding the *tprD* (*tp0131*) locus are spliced together by potentially overlapping 4-bp repeats. Out of 6,372 total *tprK* sequences, we were able to reconstruct 5,911 sequences (92.76%) using the previously defined internal repeats with over 95% coverage of the variable region, accounting for 95.0% of the total number of reads. Of the remaining 461 sequences that were unable to be reconstructed, 43.4% belonged to V1, accounting for 88.5% of the remaining number of reads. In V1, there frequently was no donor site segment spanning across the largest junction between repeats, of which V1 only has one: GCAT. Examples of these sequences, including the V1 clonal sequence in culture, are displayed in S5 Fig.

We then selected eight sequences absent in inoculum that reached ≥0.4% relative frequency to examine how these novel *tprK* sequences were being constructed from donor site segments (Fig 7). We discovered several different approaches to generating new alleles. In all cases examined, the new sequence shared at least one donor site with the highest frequency sequence (the “clonal” sequence), usually as a shorter segment than the original. Six of the eight new sequences had exactly one donor site that was not originally in the clonal sequence. Of these, two were present in the inoculum in the most common non-clonal sequence, two were present in the inoculum in one low frequency sequence, and two were not found in any other sequence in the inoculum. In addition, one new sequence in V6 was generated by splicing together segments taken from different parts of the same existing donor site (V6-DS53), or potentially by deletion of the sequence between the two segments.

**Fig 7.**
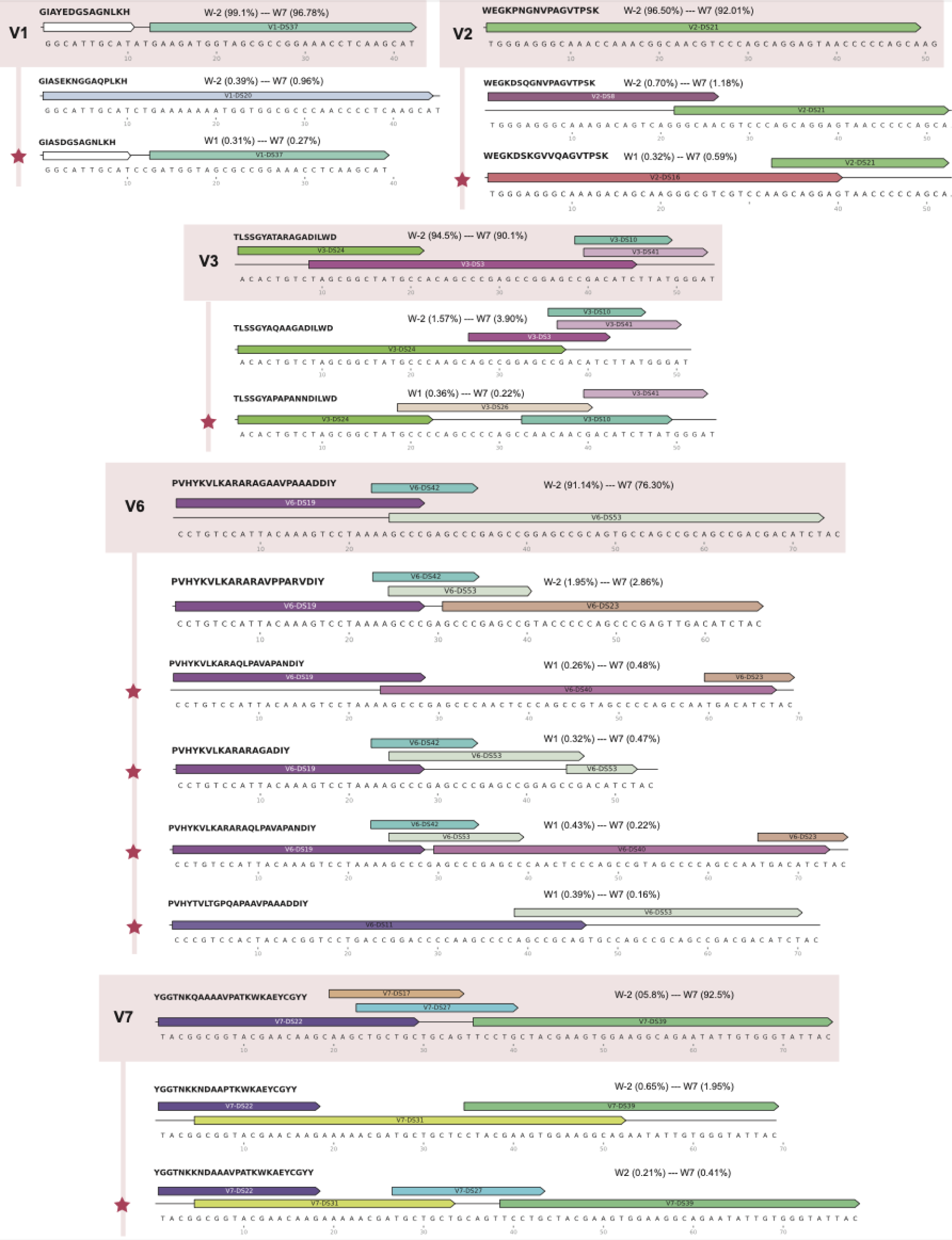
Donor site contributions for *de novo tprK* V region alleles present in any timepoint above 0.4% relative frequency. Each variable region with *de novo tprK* alleles is depicted, with the clonal sequence at the top and highlighted in salmon, followed by the next highest sequence in relative frequency, and then *de novo* alleles absent from inoculum, demarcated by red stars. Donor site segments are illustrated with color-coded arrows above their respective *tprK* sequences for each variable region. White donor site segments in V1 are segments that are present in all V1 donor sites. The translated amino acid sequences are at the top left of each allele. Range of relative frequencies of each allele are reported, with week numbers and respective relative frequencies.

We further investigated additional non-immune related causes for the greater antigenic variation in V6. One potential explanation would be that V6 allows for a greater number of donor site segments, leading to more potential for diversity. However, we found no significant difference between number of donor site segments in V6 compared to the other variable regions (S6 Fig). We additionally found no significant differences in %GC content that could indicate evidence of GC-biased gene conversion, a described non-adaptive evolutionary process selecting for higher GC content in bacteria [30, 31].

## Discussion

Here, we used next-generation sequencing to examine longitudinal evolution of TprK in multiple model systems: immunocompetent rabbits, immunosuppressed rabbits, and *in vitro* culture. Consistent with previous reports on the role of *tprK* in immune evasion [16,28,32,33], we found significantly greater *tprK* diversity in V6 during syphilis infection in treponemal samples obtained from immunocompetent rabbits compared to immunosuppressed ones, and in an immunocompetent rabbit compared to culture. Interestingly, however, we also observed that in treponemes from both immunosuppressed rabbits and in culture, *tprK* accumulates significantly more V6 variants compared to other variable regions, suggesting that immune selection is not the sole mechanism for diversity accumulation in V6. Given the reproducibility of *tprK* sequences across our two culture replicates and their tendency towards more consistent identity over time, this increased generation of TprK heterogeneity in V6 in the absence of immune pressure is unlikely to be a purely stochastic process.

Thus far multiple studies have shown that in, rabbit models, there is an active accumulation of variable TprK alleles over the course of infection, which may be attributable to selection by the host immune response [16, 34]. At minimum, five of the seven variable regions (V2, V4-V7) elicit variant-specific antibody responses [35]. Our data confirmed significantly greater diversity in V6 associated with presence of immune pressure, when comparing immunocompetent and immunosuppressed rabbits. Previous experiments on these same rabbits used a combination of fragment length analysis (FLA) and Sanger sequencing of ∼10 clones per *tprK* amplicon to estimate diversity [16]. Sanger sequencing results showed more rapid accumulation of sequence diversity in V6, while FLA also detected increased diversity in V4, V5, and V7. In this study, our deep sequencing approach mainly detected a significant difference in accumulation in V6, recapitulating the previously observed decreased sensitivity of sequencing compared to FLA [16]. This difference in detection may be due to the limited copy numbers in the later weeks, particularly for immunocompetent rabbits, which would falsely lower diversity estimates, or could be due to our stringent method of calling high-confidence sequences via presence in two library preparations. When we examined cultured treponemes, a similar and significantly increased accumulation of diversity in V6 was seen in a control rabbit compared to the cultured cells, as measured both by Pielou’s evenness score and number of unique sequences. In this experiment, we did observe a significant accumulation of diversity arise in other variable regions (V2, V3, V4, and V7) in the control rabbit compared to in culture. This could be attributed to the low number of replicates in our *in vitro* vs *in vivo* experiment, with only one rabbit and two sets of culture, or alternative sampling of lesions in this rabbit. Overall, these results highlight the association of immune pressure with increased antigenic variation.

We further showed that immune pressure is not strictly necessary for increased sequence variation in V6. In both immunosuppressed rabbits and in culture, V6 has higher rates of accumulation of unique sequences compared to the other variable regions. Further research is needed to determine the exact mechanism of this increased variation. We found no significant differences in number of donor site segments in V6 when using previously characterized internal repeats and known *tprK* sequence anatomy [13]. This could potentially be due to lack of our recognition of <4 bp donor site segments due to limitations in blastn settings. We additionally found no significant differences in %GC in V6 that would raise the possibility of GC-Biased Gene Conversion (gBGC), a non-adaptive evolutionary process selecting for higher GC content in bacteria [30, 31].

Our study was also limited by the only near-clonality of our *T. pallidum* inoculum. In our initial infecting isolates we observed a greater number of sequences in V6 as compared to other variable regions (S2 Table). As a result, it is difficult to fully eliminate the possibility that this greater diversity from the onset allowed for the increased rate of accumulation of *tprK* sequence heterogeneity in V6, especially if variable region diversity generation is a non-linear process. In future studies, using limiting dilution to obtain true clonality would better inform our understanding of *T. pallidum* generation of antigenic variation. These procedures are being tested and established in the *in vitro* culture model.

The generation of novel TprK variants may not purely be due to random chance. We discovered remarkably consistent relative frequencies of V region alleles across our two replicate sets passaged in culture (r^2^ = 0.9998). The maximum relative frequency of an allele present in one culture set but not the other at the same timepoint was 0.62%. In addition, over time, relative frequencies—including initially diverging ones—became closer in identity, and often evolved in tandem. This brings up several different possibilities, including potential adaptation to culture and the existence of a preset stepwise mechanism of antigenic variation generation. Further replicates with more strains that are passaged over a longer period will be required to more robustly demonstrate the degree of stochasticity in the basal generation of *tprK* variable region alleles.

We further examined novel *tprK* variable region sequences generated in culture that were functionally absent in the inoculum, of which there were a total of 128. These new *tprK* sequences stayed at low frequencies over time, reaching a maximum of 0.79% relative frequency. While all these sequences were present at <0.1% relative frequency in the inoculum, a majority were only in one library preparation. Given the extremely low counts of these sequences, their presence in the inoculum could be due to low-level index hopping or low-level polymerase error. We considered these sequences as new and appropriate for further analysis to better understand how *T. pallidum* generates novel sequences. Using previously characterized *tprK* anatomy involving non-reciprocal segmented gene conversion of donor site segments from the *tprD* locus spliced together with internal 4 bp repeats, we explored several different sequence structures. For most cases *in vitro*, sequence is built from the pre-existing donor sites in the clonal sequence. However, several new *tprK* alleles drew from donor sites in other lower frequency sequences, and still others were built from donor sites not present in any other sequence in the inoculum. Further refinement of this mechanism is needed for firmer conclusions. In addition, many sequences, including the V1 clonal sequence *in vitro*, are unable to be constructed with the previously described internal repeats [13]. More investigation on *tprK* anatomy and potentially addition of new internal repeats is needed to account for these sequences. Understanding how these new sequences are being generated would better inform *T. pallidum* pathogenesis and potentially allow us to develop targeted drugs interfering in this process. Finally, our demonstration of longitudinal *tprK* evolution during *in vitro* culture allows for potential interrogation of mechanisms of gene conversion, especially given the new availability of genetic engineering approaches in *T. pallidum* [8].

## Funding information

This work was supported by NIAID U19AI144133. The funders had no role in study design, data collection and analysis, decision to publish, or preparation of the manuscript.

## Competing interests

ALG receives funding for central testing from Abbott and research grant support from Merck and Gilead, unrelated to the work performed here.

## Supporting Information

S1 Table. Sequencing primers used in this study.

S2 Table. Number of variable region sequences and diversity measures for the 7 variable regions of TprK for all samples analyzed in this study.

S3 Table. Mean weekly decrease of most abundant sequence passaged in immunosuppressed rabbits and *in vitro* culture by variable region.

**S1 Fig.**
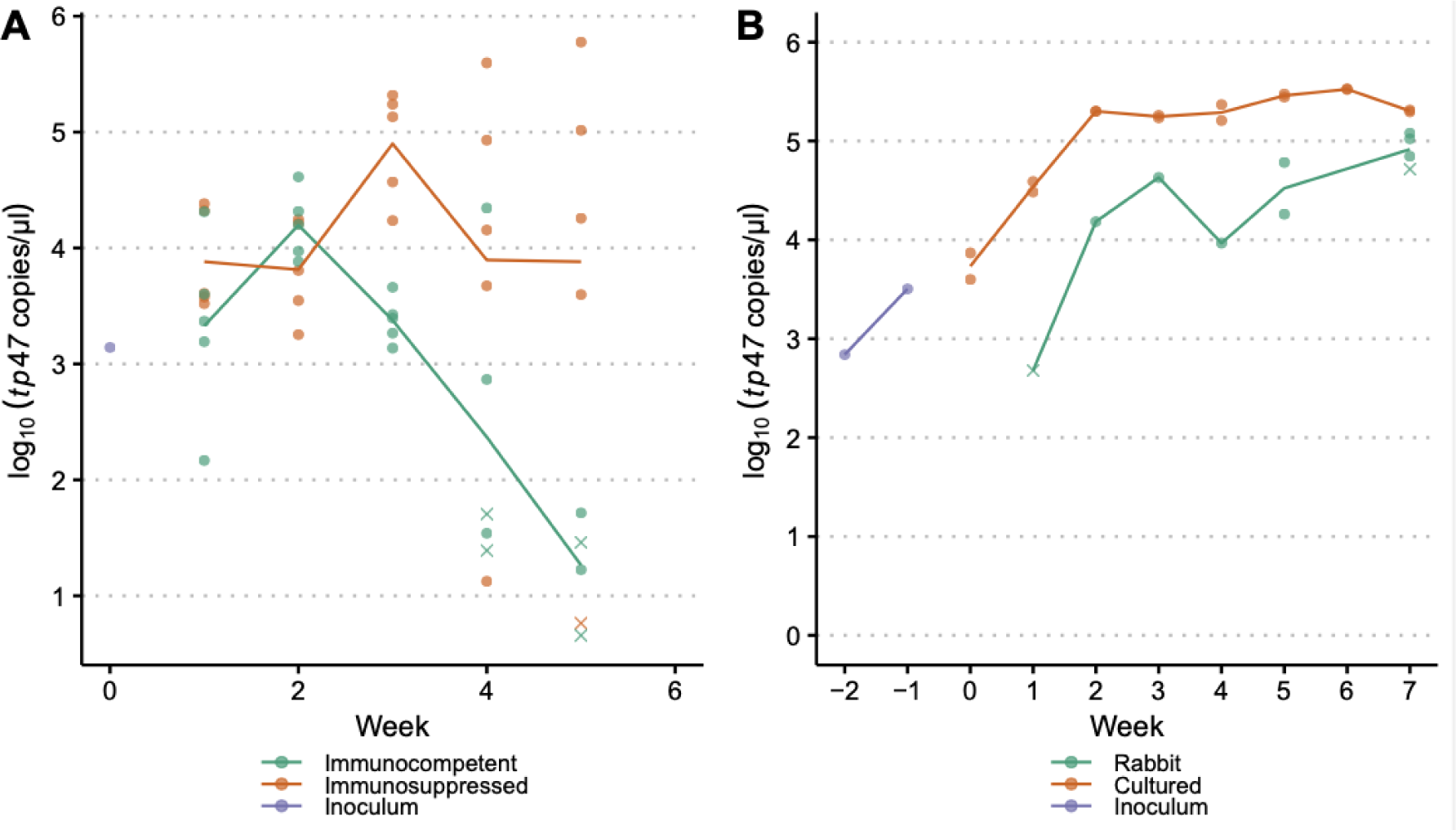
Treponemal load of samples passaged in (A) immunocompetent vs. immunosuppressed rabbits and (B) immunocompetent rabbit vs. culture. Each unique sample is represented by either a dot or a cross, where a cross indicates that we were unable to successfully recover sequence from the sample. Lines show the mean treponemal load, as quantified by the log of *tp47/*μl, and are grouped and color-coded by passage type.

**S2 Fig.**
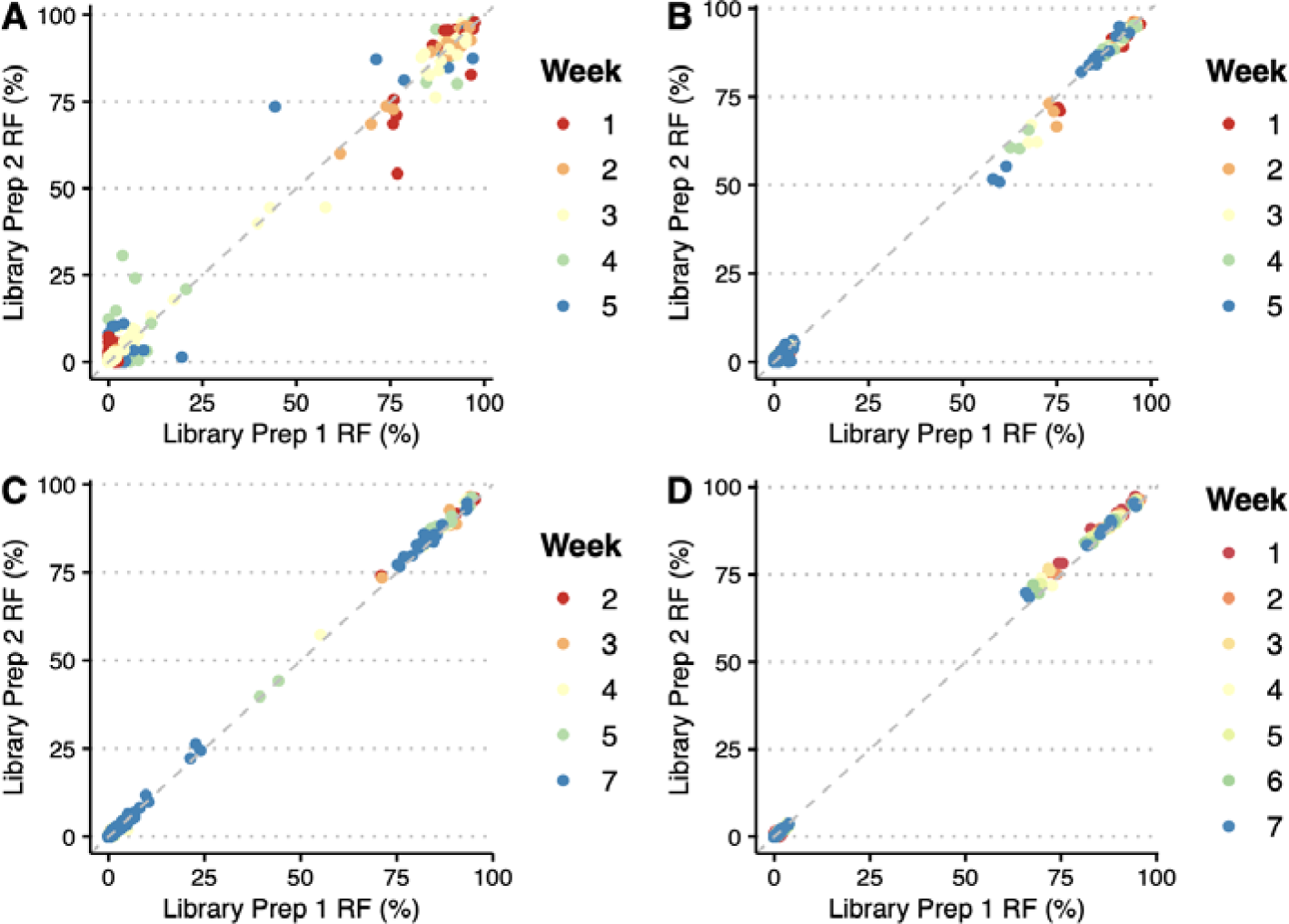
Reproducibility of technical replicates in two Illumina library preparations across samples passaged from (A) immunocompetent and (B) immunosuppressed rabbits in immunocompetent vs. immunosuppressed rabbits experiment, and (C) immunocompetent rabbit and (D) culture in *in vivo* vs. *in vitro* experiment.

S3 Fig. Longitudinal tprK allele frequencies for immunocompetent vs. immunosuppressed rabbits. Only *tprK* alleles existing across multiple collection timepoints are present. Alleles that have >10% frequency at any time point are specifically labeled at the top of each variable region.

S4 Fig. Longitudinal tprK allele frequencies for *in vivo* vs. *in vitro* experiment. Only *tprK* alleles existing across multiple collection timepoints are present. Alleles that have >10% frequency at any time point are specifically labeled at the top of each variable region.

**S5 Fig.**
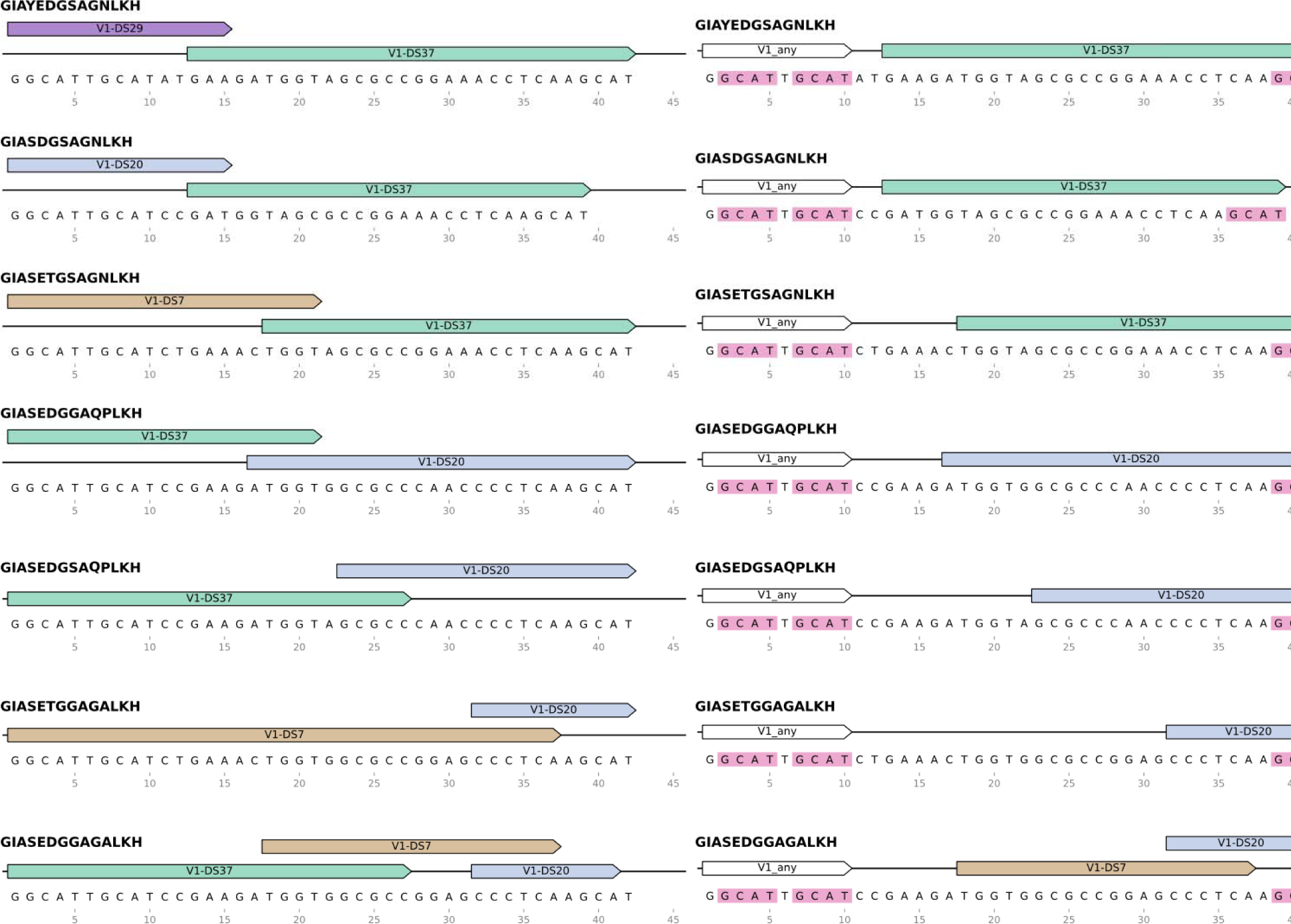
Construction of V1 sequences with and without accounting for internal repeats. On the left are sequences constructed with the longest possible donor site segments. Paired on the right are the same sequences following the internal repeat structure rules. Donor sites are represented as labeled color-coded arrows. The first segment of the sequences on the right can be represented by any known V1 donor site. GCAT repeats are highlighted in pink.

**S6 Fig.**
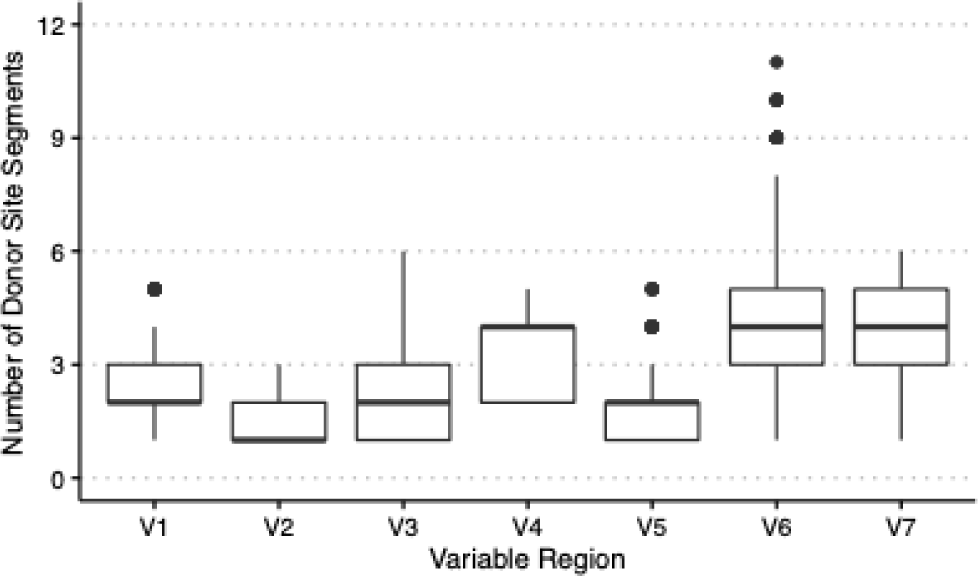
Number of donor site segments in donor site recombination possibilities for each variable region. Internal 4-bp repeats are not included in the number of donor site segments. All combination possibilities with the highest % coverage were kept. Outliers are shown as dots outside of the boundaries of each boxplot.

