## Supplementary figures and images for "Longitudinal accumulation of *in vivo* and *in vitro-*grown *Treponema pallidum* subsp. *pallidum* TprK variants in the presence and absence of immune pressure"

### Fig S3

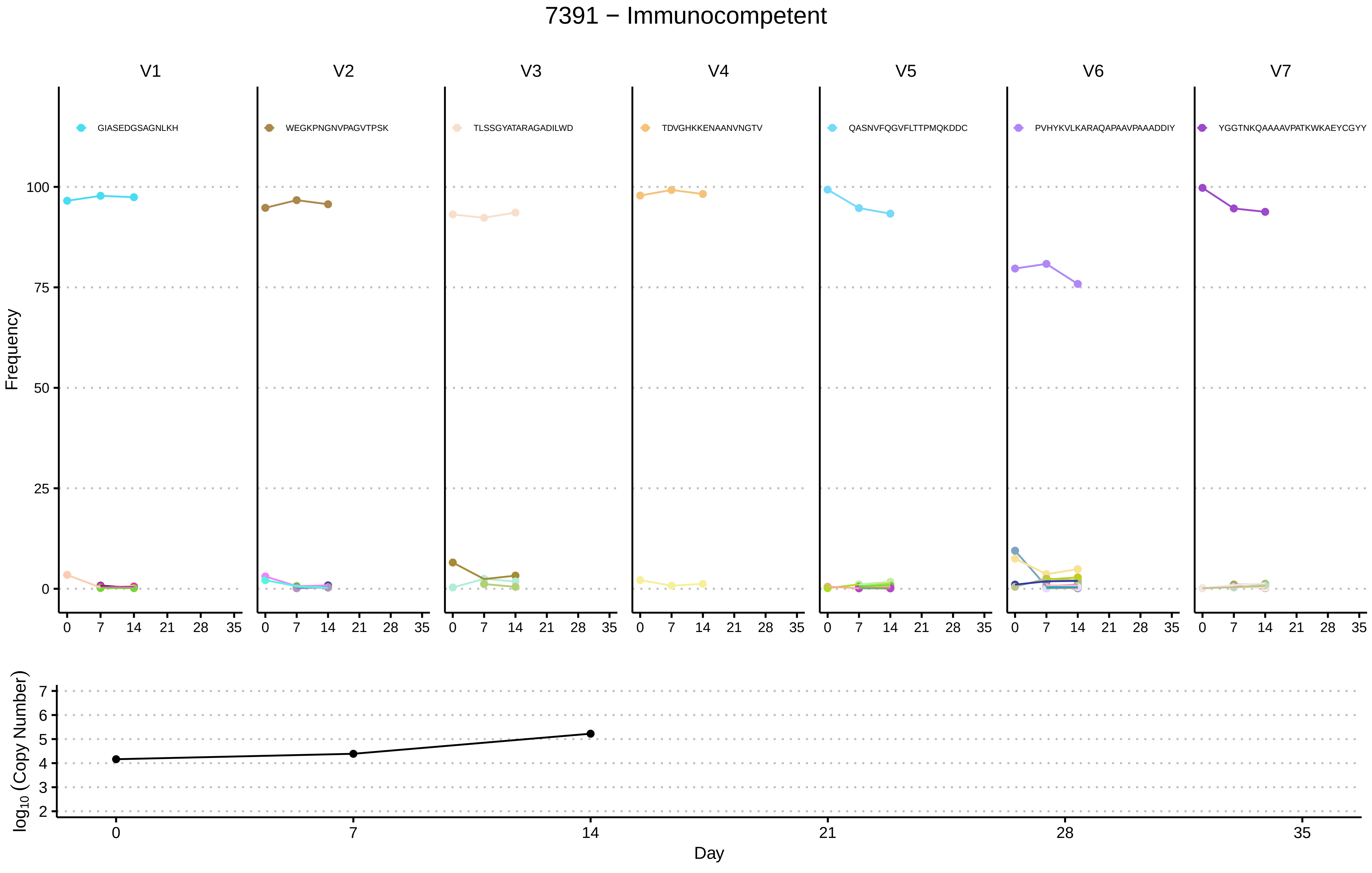

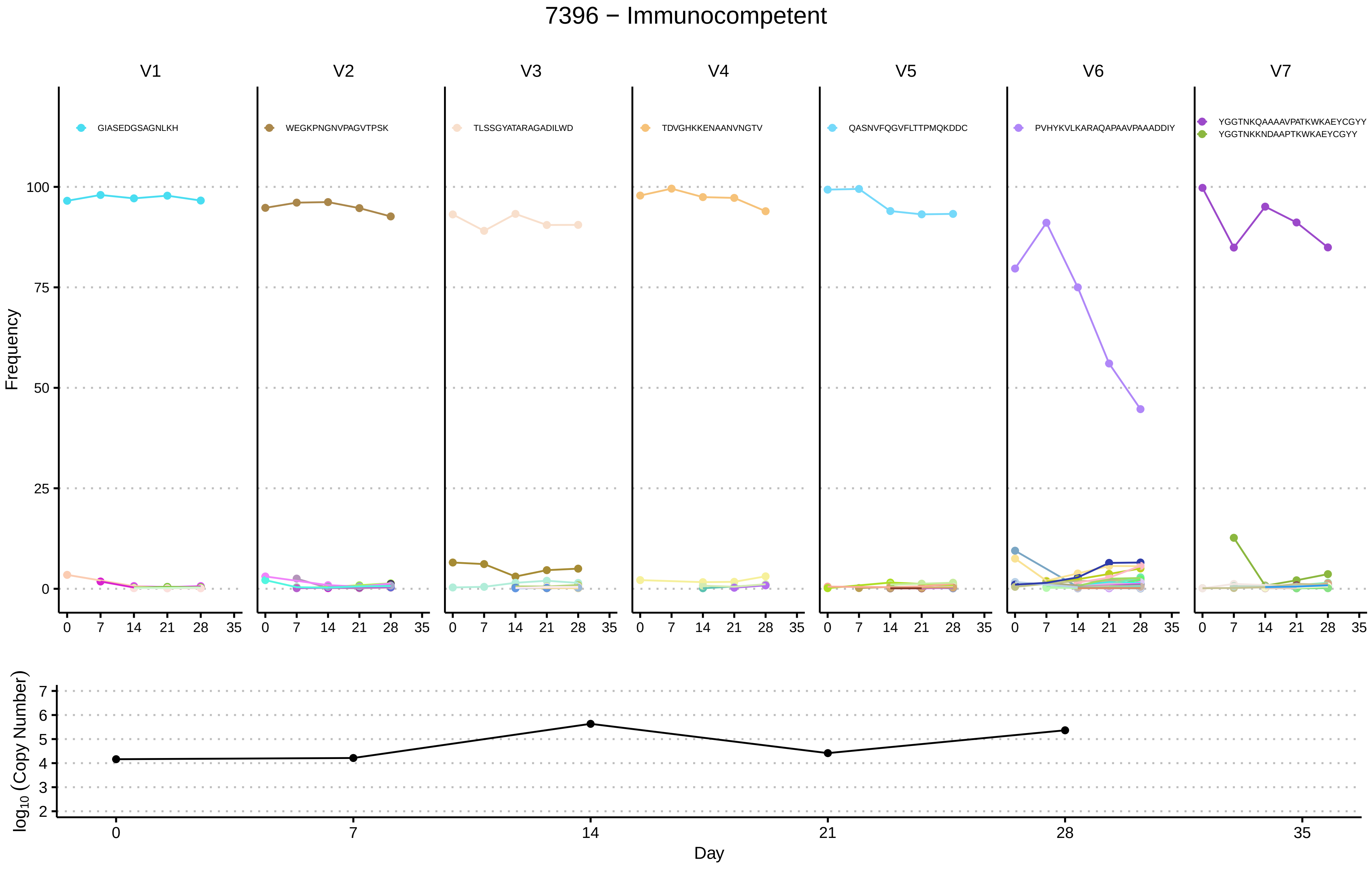

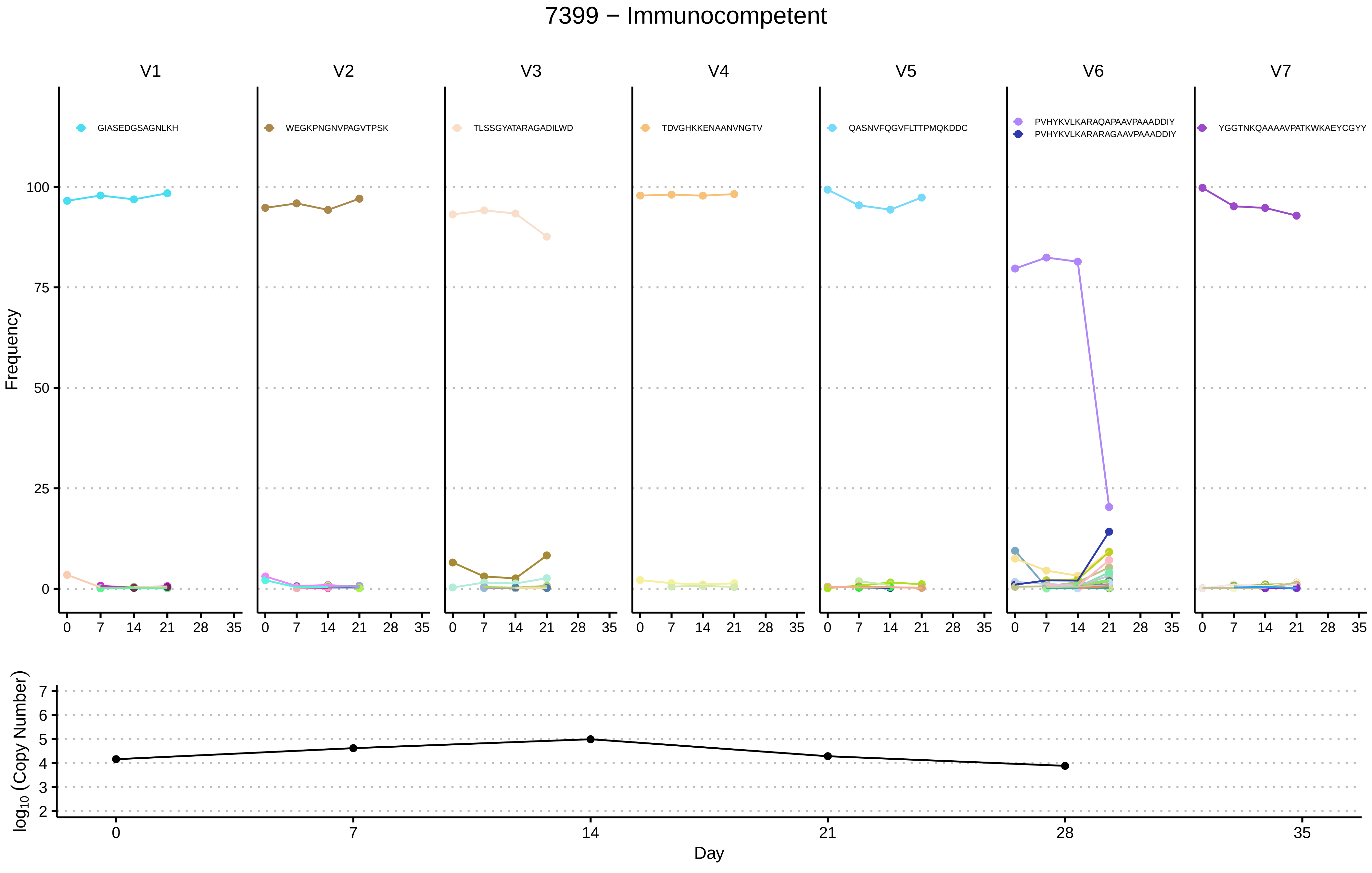

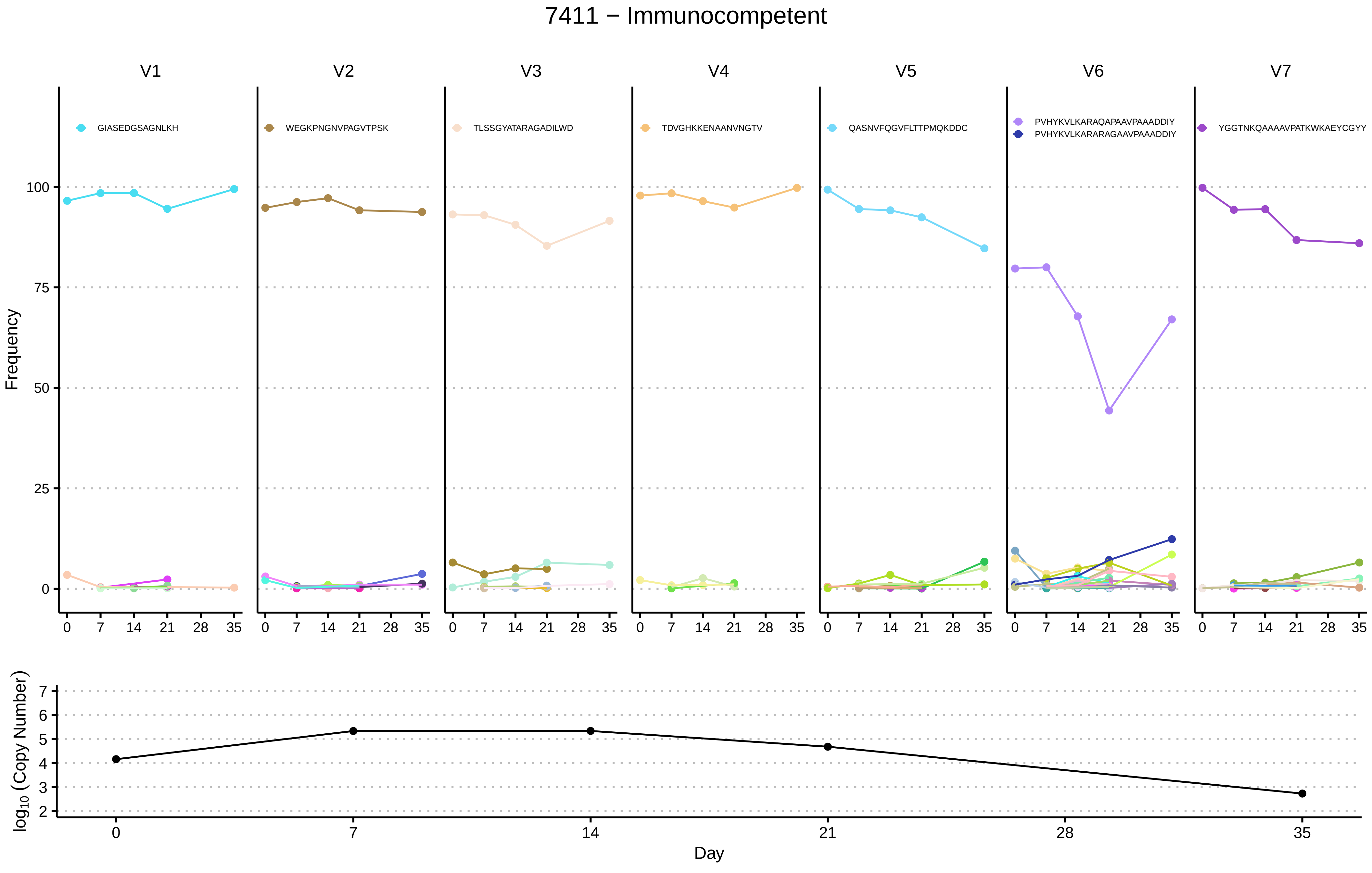

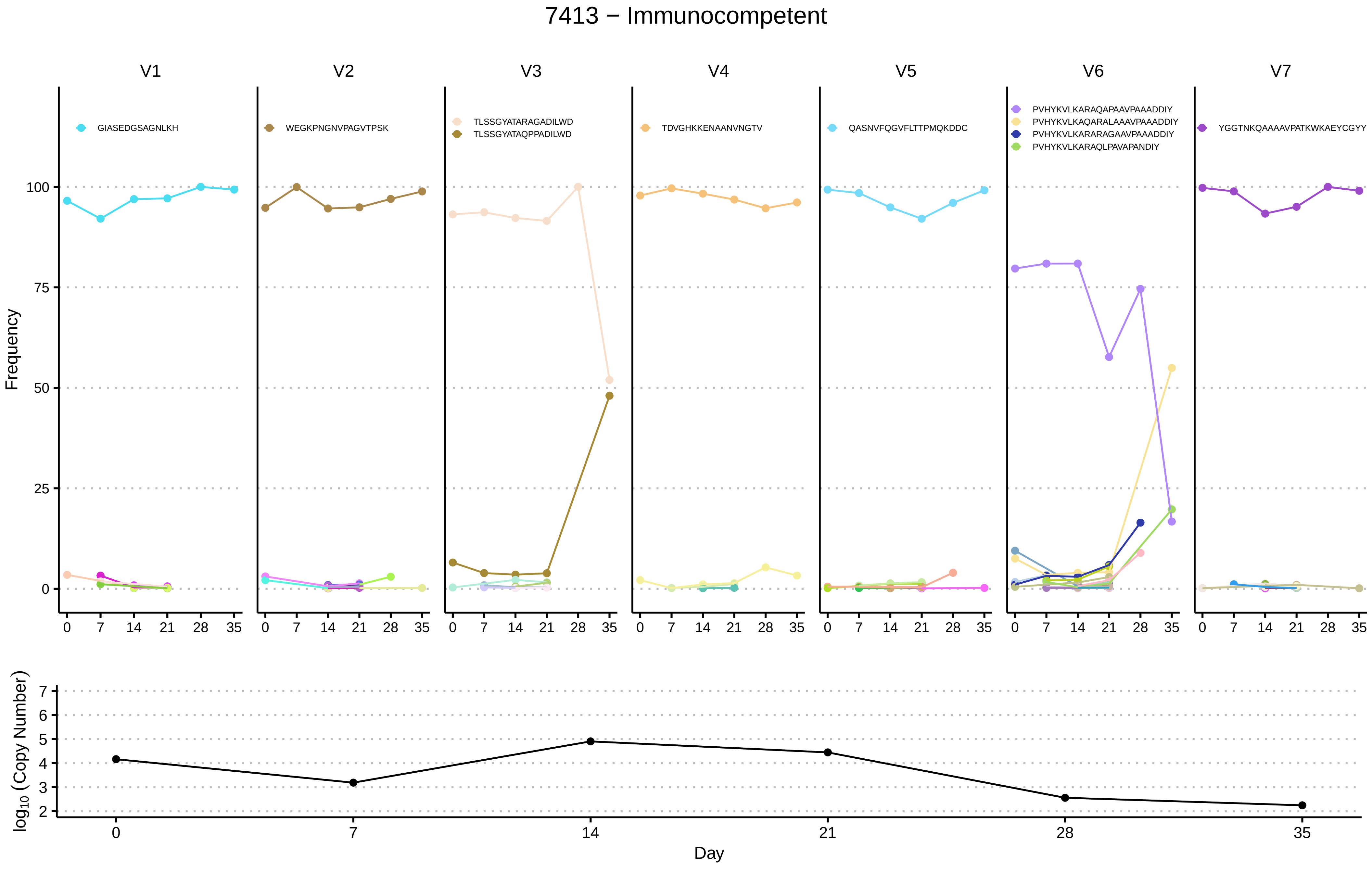

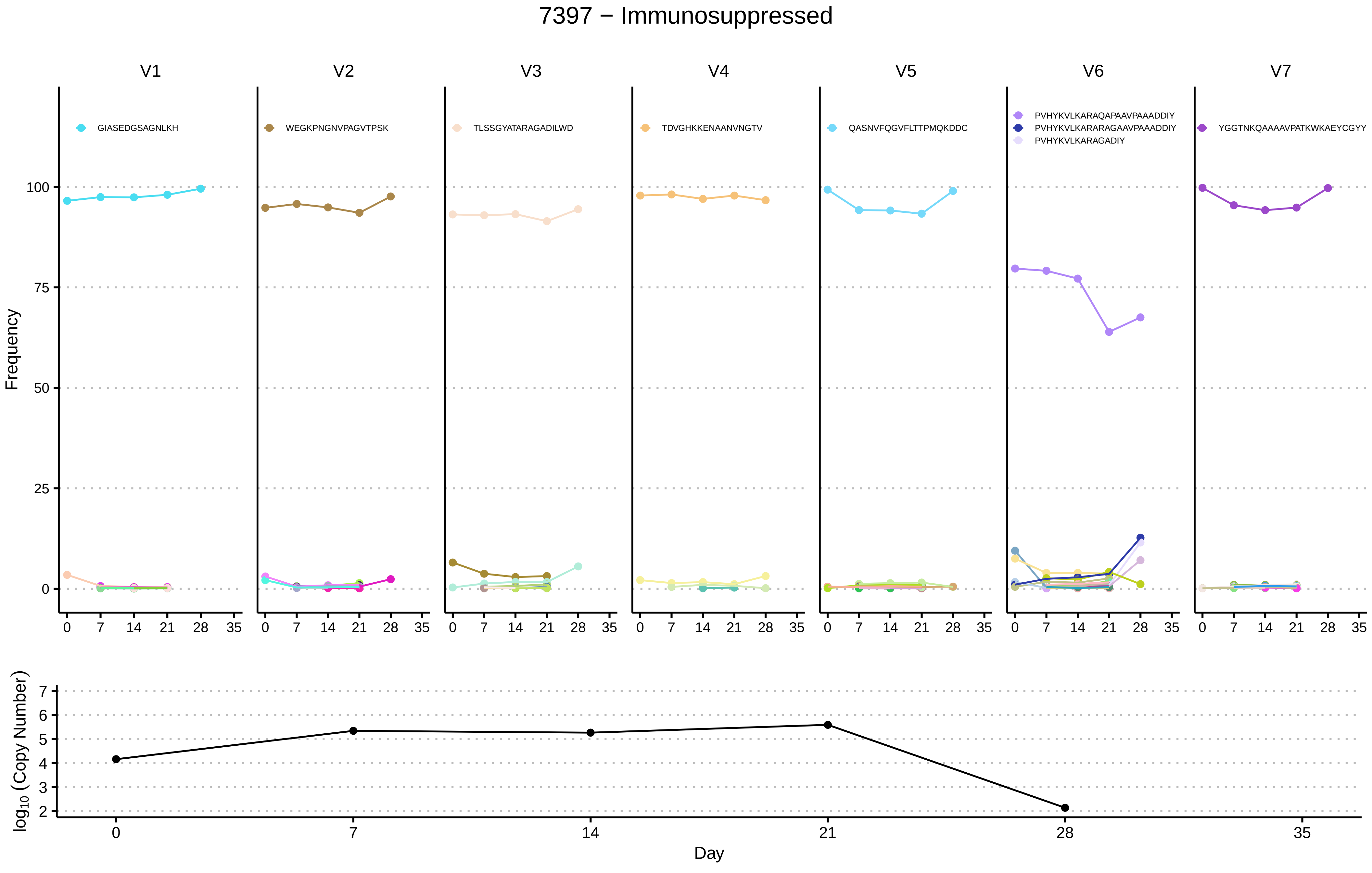

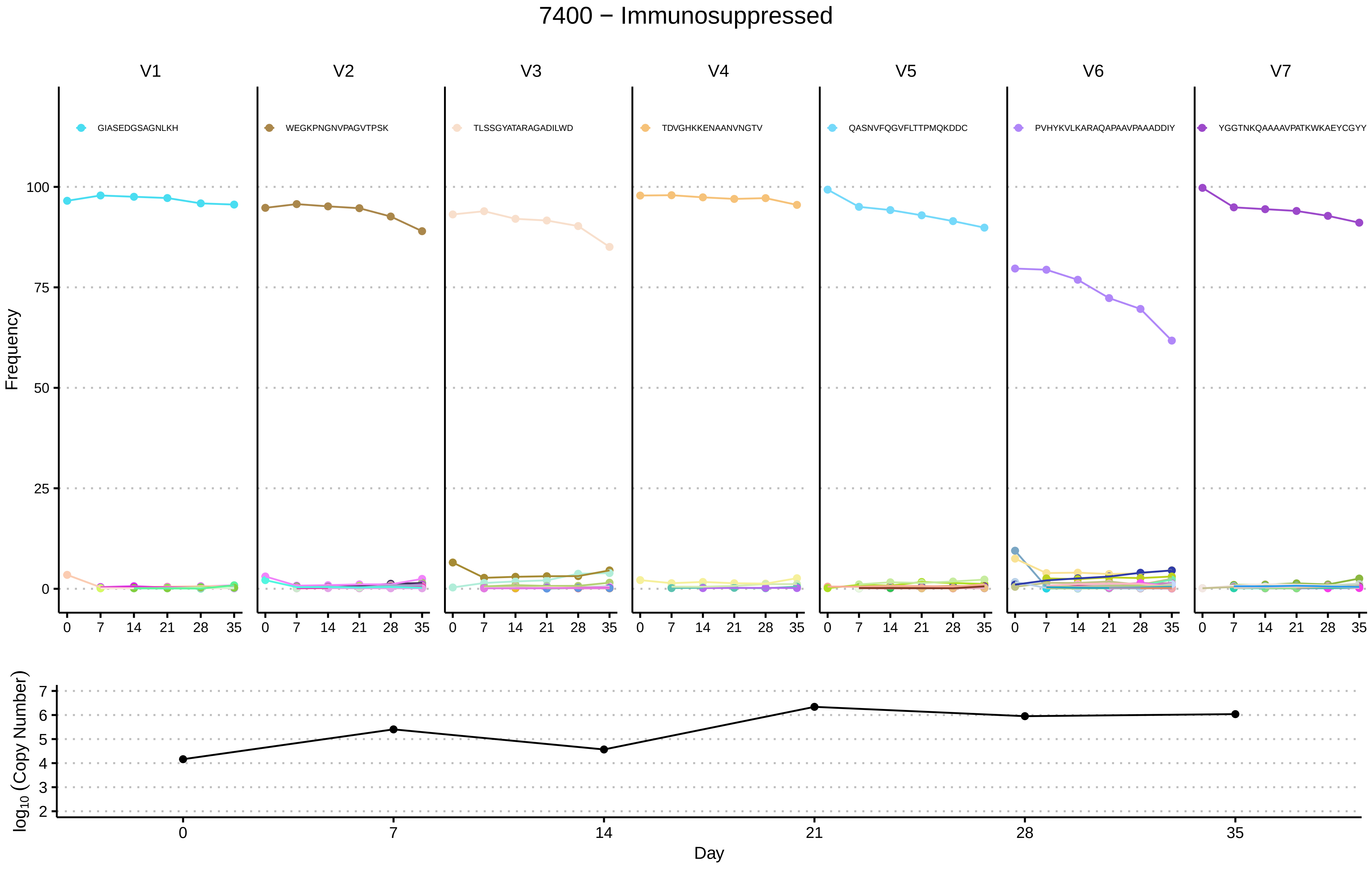

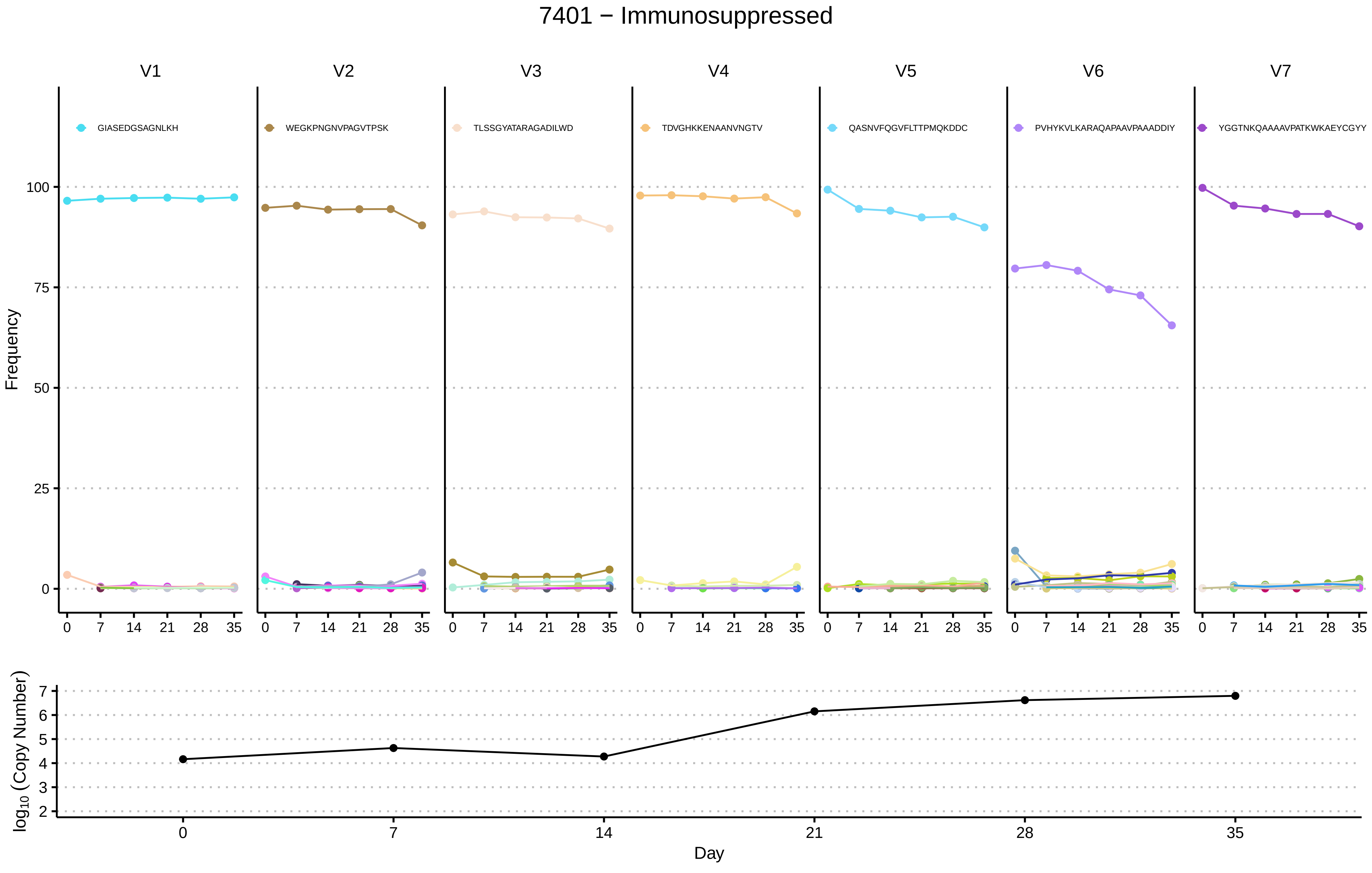

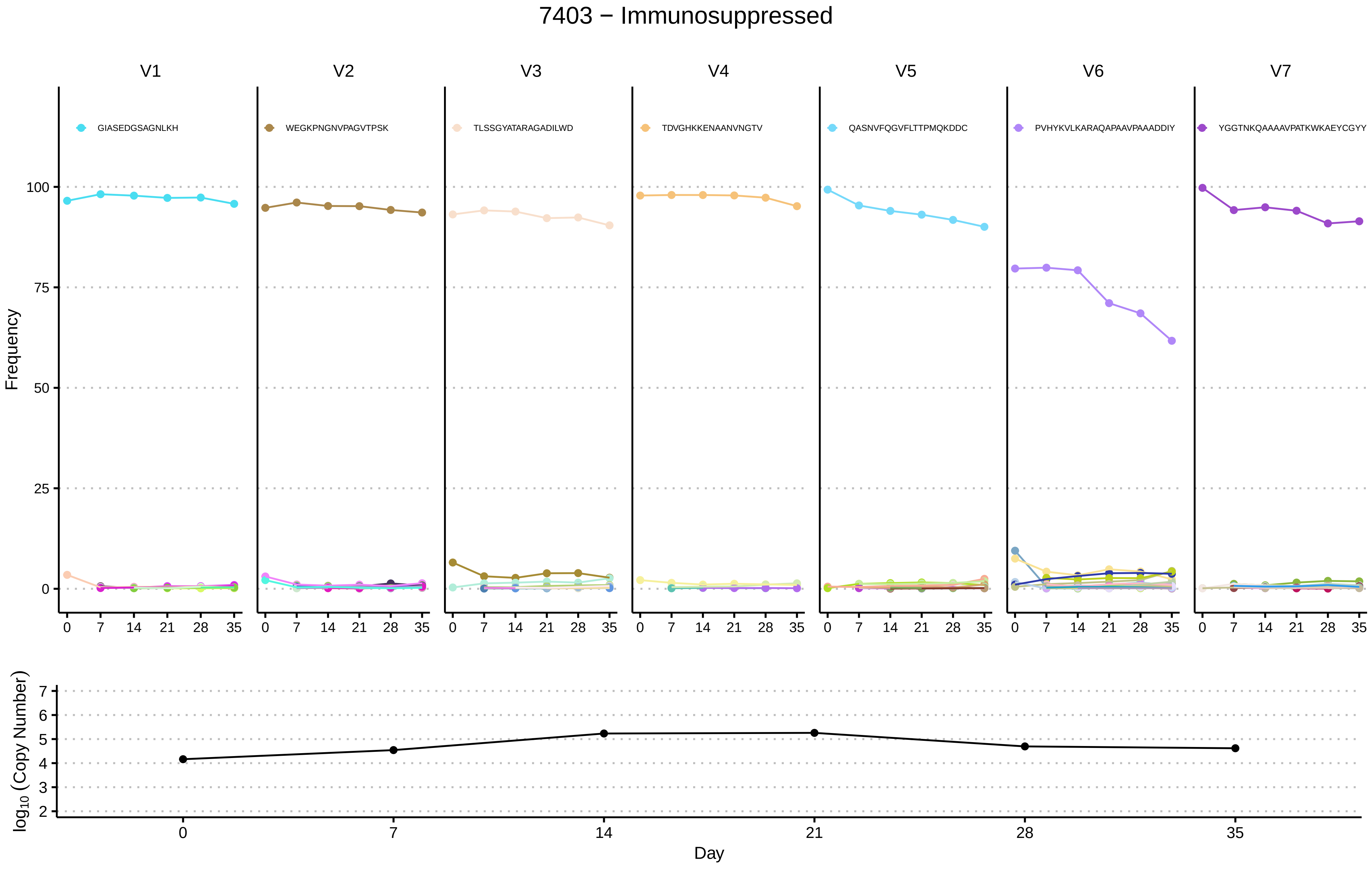

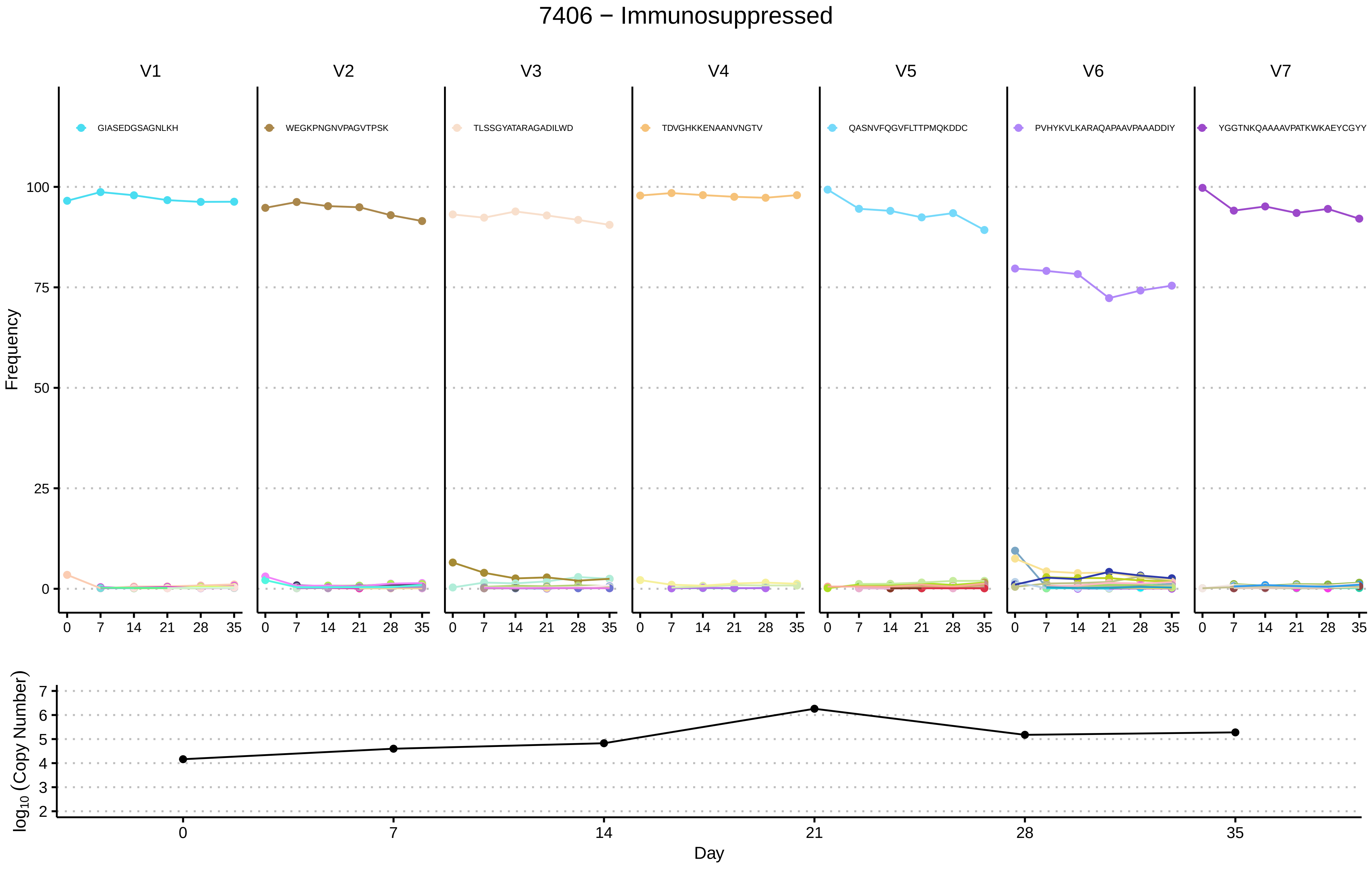

### Fig S4

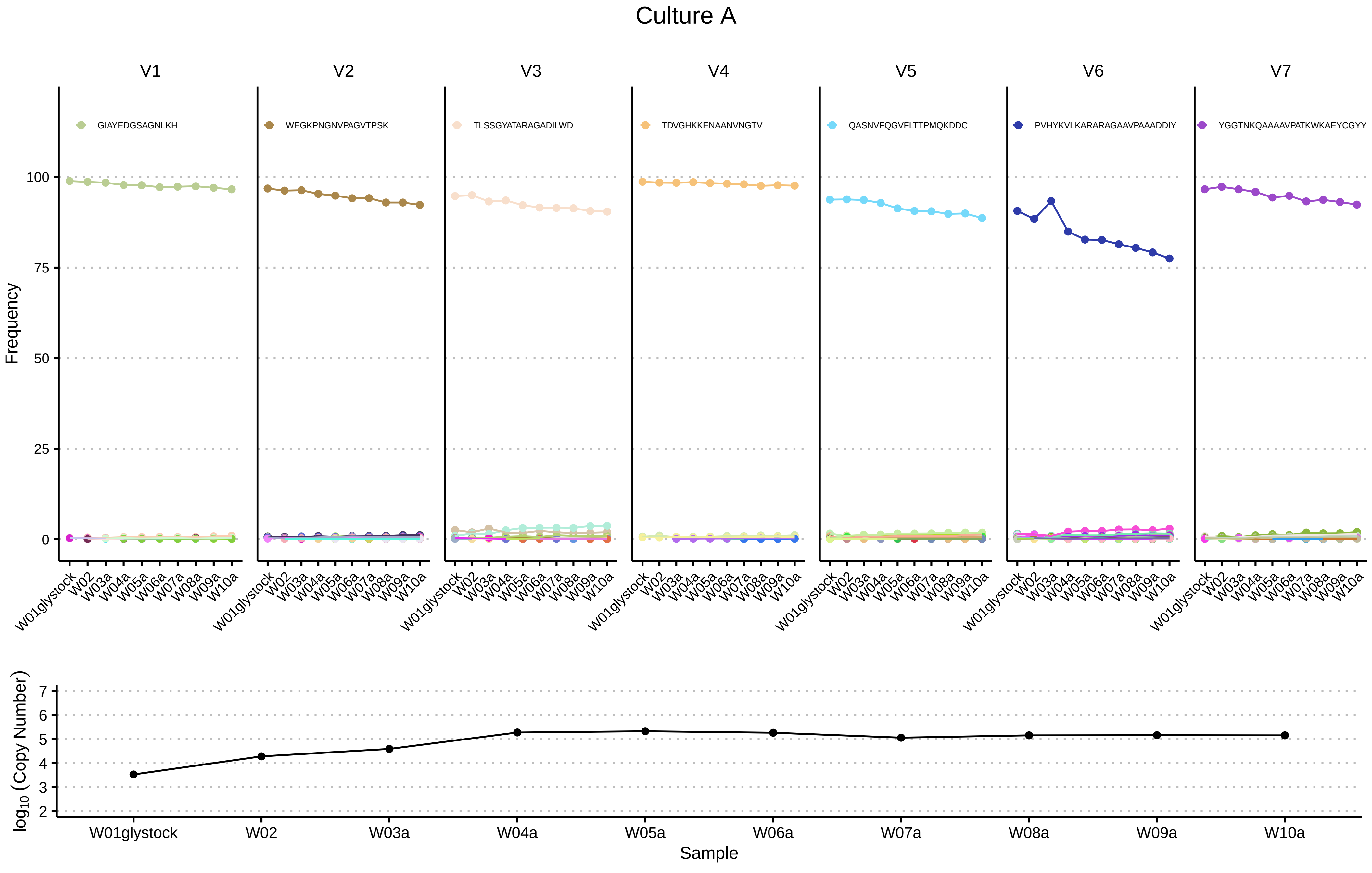

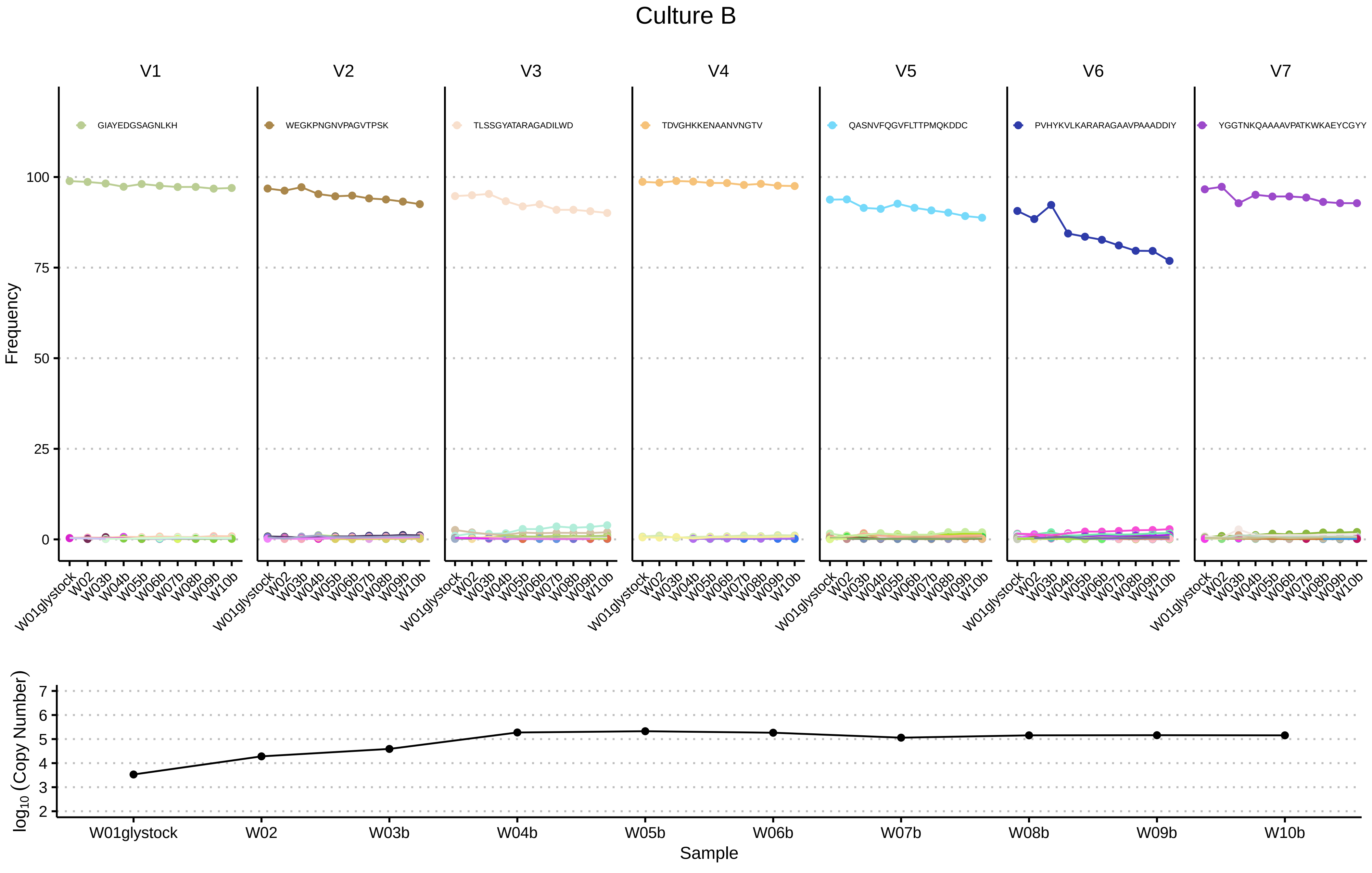

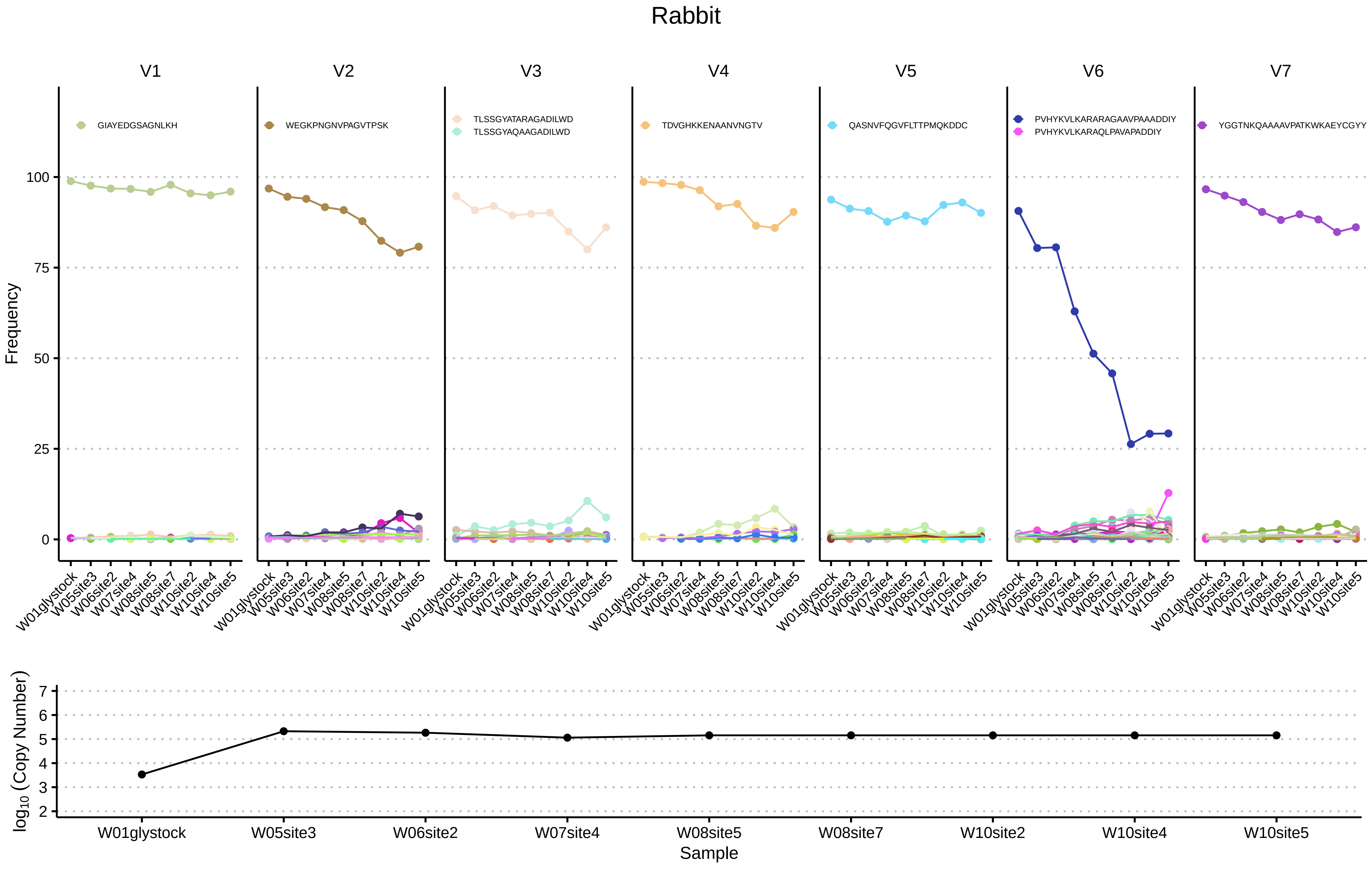
